# An integrated systems biology approach establishes arginine biosynthesis as a metabolic weakness in *Candida albicans* during host infection

**DOI:** 10.1101/2025.01.11.632533

**Authors:** Shuvechha Chakraborty, Indumathi Palanikumar, Yash Gune, K.V. Venkatesh, Karthik Raman, Susan Idicula-Thomas

## Abstract

*Candida albicans* (CAL), one of the leading causes of fungal infections affecting nearly 70% of the population, poses a significant global health threat. With the emergence of drug-resistant strains, mortality rates have reached a staggering 63.6% in severe cases, complicating treatment options and demanding the discovery of novel therapeutic targets. To address this pressing need, we employed a unique multidisciplinary approach to elucidate the metabolic pathways that enable CAL to switch from a commensal to a virulent state. Condition-specific genome-scale metabolic models (GSMMs), along with a novel integrated host-CAL model developed in this study, highlighted the central role of arginine (Arg) metabolism and uncovered *ALT1*, an arginine biosynthesis enzyme, as a critical metabolic vulnerability in CAL virulence. Heightened expression of arginine biosynthesis genes indicated that increased arginine synthesis mainly occurs through proline intermediates during host interaction. Significantly impaired virulence and *in vivo* pathogenicity of *ALT1*-deleted CAL highlighted the potential of targeting arginine metabolism as a novel strategy to combat antifungal resistance and underscored the power of integrating systems biology with experimental approaches in identifying new therapeutic targets.

**Graphical abstract:** 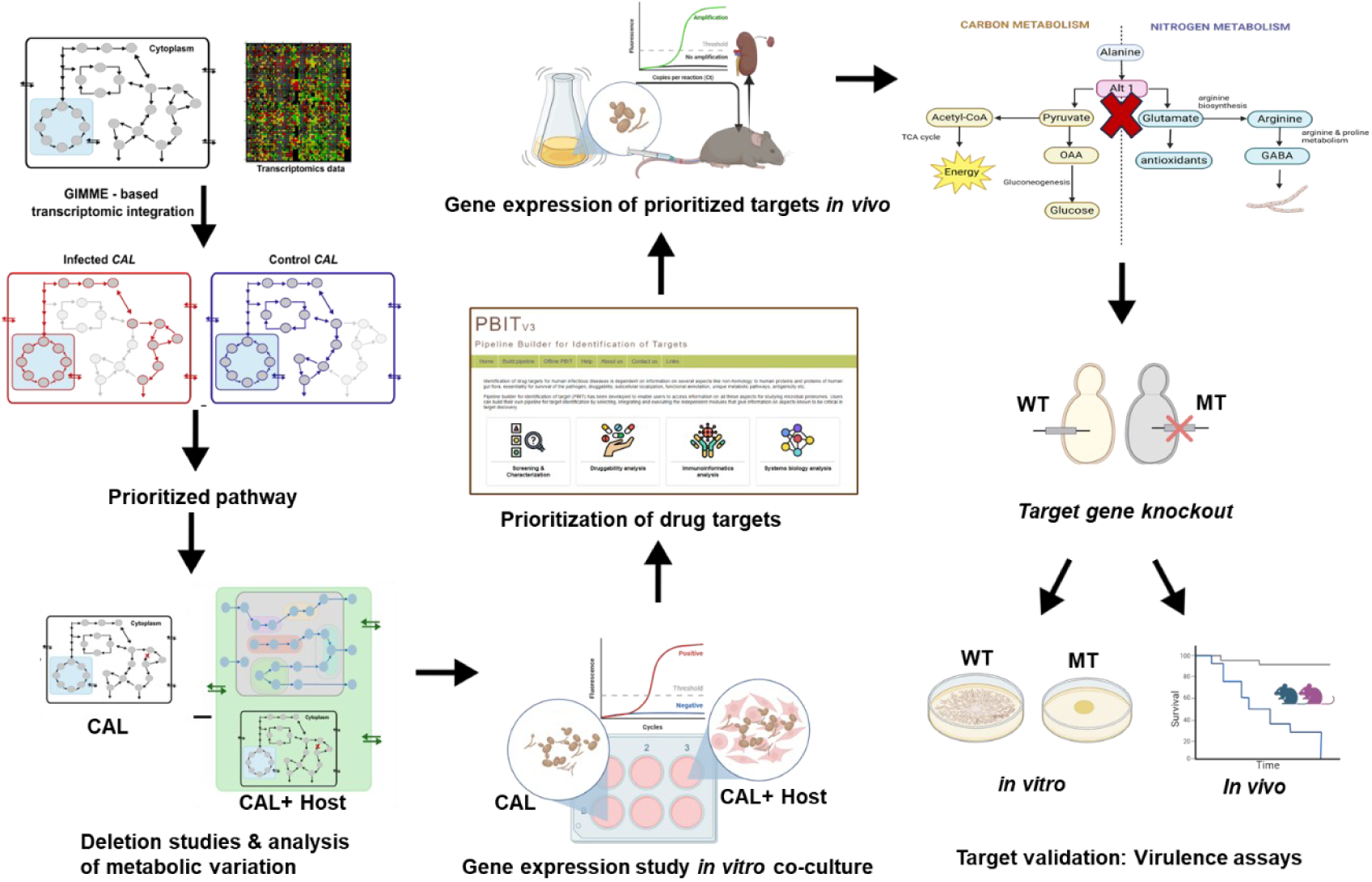

## 1. Introduction

Candidemia—the presence of Candida species in the bloodstream—is a leading healthcare-associated bloodstream infection^1,2^. It is the most frequently recognized condition linked to invasive/systemic candidiasis^3^. The incidence of systemic infections caused by CAL and non- *albicans Candida* species (NACs) is alarming, with over 150 million cases reported annually and a staggering mortality rate of 63.6%^4^. Although NACs are increasingly becoming a concern, CAL remains the predominant species, isolated in over 50% of reported *Candida* infections^5,6^.

Current antifungal treatments focus on targeting the fungal cytoskeleton, specifically the cell wall and plasma membrane, through four major classes: azoles, echinocandins, polyenes, and allylamines. However, the rampant overuse of these antifungals in clinical settings and intensive care units has accelerated the emergence of drug-resistant *Candida* species. In immunocompromised patients, existing treatments are failing to deliver adequate protection, consequently leading to rising mortality rates in these vulnerable populations^7^. Despite the escalating threat posed by drug resistance, the development of new antifungal treatments has not kept pace with *Candida’s* rapid evolution. This urgent situation has led the World Health Organization (WHO) to designate *Candida* species as critical and high-priority on its 2022 fungal priority pathogens list^8^.

*Candida* is an opportunistic pathogen that effectively thrives in host environments often characterized by low nutrients and oxygen due to its remarkable metabolic flexibility. Environmental cues such as nutrient scarcity, hypoxia, pH fluctuations, and temperature shifts decisively drive the transformation between its yeast form and more virulent hyphal phenotypes. The metabolic pathways of *Candida* are critical in mediating its immune responses, pathogenicity, and drug resistance^9,10^, such as dynamic changes in glucose metabolism significantly influence adhesion, stress resistance, and immune invasion in animal models^11^. Consequently, these metabolic pathways present compelling targets for the advancement of innovative therapeutic interventions^12,13^.

Conventional drug target identification is a labor-intensive and costly endeavor, even amid advancements in biotechnology. While computational approaches have been used for decades, recent innovations in constraint-based modeling and the availability of high-quality metabolic models have revolutionized this process, enabling the effective shortlisting of metabolic targets in diseased conditions^14^.

Genome-scale metabolic models (GSMMs) have emerged as valuable tools for understanding the metabolic behaviour of organisms^15,16^. By integrating computational models and experimental data, these models accurately map the metabolic state of the organism. As a result, GSMMs can be effectively used to simulate metabolic fluxes under various conditions^17,18^ and have consistently demonstrated their capability to predict drug targets for numerous bacterial and eukaryotic pathogens^19,20^. Recently, various studies showcased the ability of GSMMs of multiple organisms (*C. parapsilosis* (iDC1003), CAL (iRV781), *C. glabrata* (iNX804) and *C. tropicalis* (iCT646)) in identifying essential metabolic entities as drug targets^21–24^.

Overlaying gene expression data onto the metabolic framework of a GSMM can enhance the representation of an organism’s functional state. Previous research^25,26^ demonstrated the effective application of a GSMM for *Mycobacterium tuberculosis* and *Plasmodium falciparum*, in genomics and metabolomics integration to assess how genetic variations influence pathogenicity and essentiality. Despite its success with *M. tuberculosis* and *P. falciparum*, this methodology remains unexplored for *Candida* spp.

In this work, we employed a multifaceted approach, combining systems biology with experimental validation, to elucidate the metabolic underpinnings of CAL virulence. We developed context-specific GSMMs to depict the metabolic states of CAL in both infected and uninfected settings and also built an integrated host-CAL metabolic model to delve into the critical role of reactions associated with the identified pathway. Detailed analysis of these models showcased the importance of arginine (Arg) metabolism in CAL pathogenicity during host interactions. Through gene expression studies with CAL co-cultured with the vaginal epithelial A-431 cell line, as well as in a murine model of systemic candidiasis, we validated the participation of Arg metabolism genes in host-pathogen interactions. A gene deletion mutant strain for CAL (MT) was created for a novel target gene *ALT1* identified from systems biology, *in vitro* and *in vivo* models. Mice injected with MT displayed a survival rate of 100%, starkly contrasting with the outcomes of those injected with the wild-type (WT) CAL strain. These compelling results indicate that disrupting the Arg biosynthesis pathway diminishes the pathogenic potential of the fungus, likely due to reduced Arg availability for gamma-aminobutyric acid (GABA) synthesis— an essential component for germ tube formation and filamentation (refer to graphical abstract).

## 2. Results

### 2.1. Context-specific GSMMs identify Arg metabolism as a crucial pathway in CAL infections

Context-specific metabolic models were constructed with *iRV781* (CAL) to capture the metabolic changes that accompany *Candida* infection. The condition-specific CAL models were generated by integrating transcriptomics data available in GEO from cell-line studies, including HUVEC and OKF6^27^ and murine infection models^28^ into *iRV781* using the GIMME algorithm, with a 10% expression threshold. These models were investigated to unravel the key metabolic alterations associated with CAL infection. After evaluating the growth of the context-specific models, variations in active reactions and functional fluxes between uninfected and infected conditions were analyzed. The models showed only minimal differences (∼3 reactions) in the number of active reactions and metabolites between the two conditions (Table S1). While the active reactions were largely consistent (98%), essentiality analysis revealed significant differences in the reactions observed, especially related to amino acid and fatty acid metabolic pathways, between uninfected and infected states. Arg biosynthesis enrichment was observed through the analysis of maximum flux and flux span reactions, highlighting its importance for survival during infection (Figures 1A and 1B) Reaction profiling of CAL at different time points (5 h and 8 h) revealed consistent activation of reactions related to Arg synthesis and atrazine degradation in both CAL-infected HUVEC and OKF6 cell lines (Table S2). Additionally, reactions related to amino acid metabolism (Histidine and Tryptophan) (Figure 1B) and fatty acid metabolism (Figure 1B) were also observed to be functionally critical and enriched in the murine systemic candidiasis model. CAL infections in both HUVEC and murine models led to increased purine, nicotinate, and glycerolipid metabolism.

**Figure 1.**
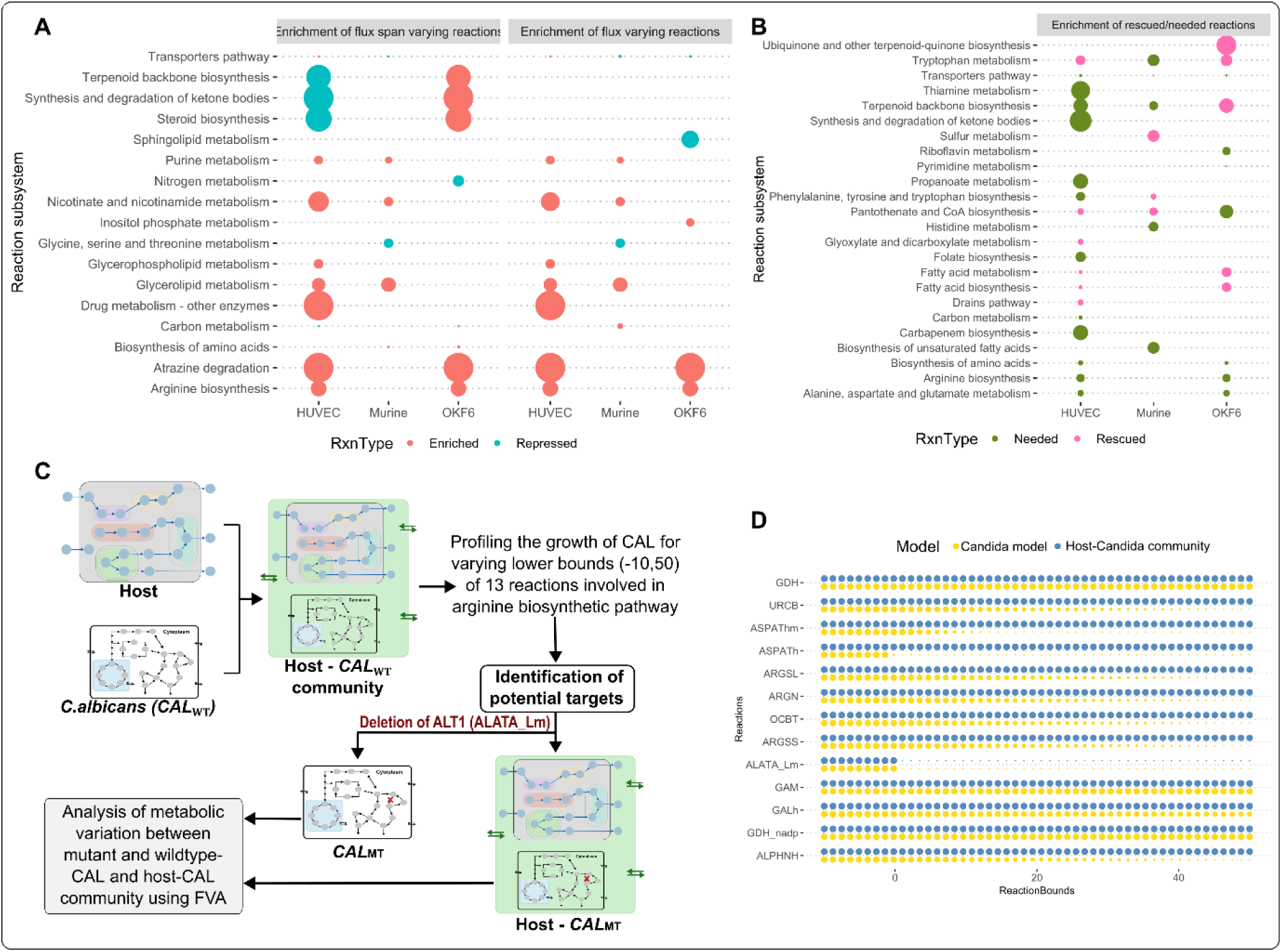
Overview of the metabolic alterations during CAL infection deciphered from flux analysis of context-specific CAL (iRV781) model (A and B) and growth profile of CAL in solitude and in host-CAL model upon metabolic flux variation of Arg biosynthetic reactions (C and D). (A) Bubble plot illustrates the enrichment or repression of the reaction subsystems in CAL during Candida infection using the maximum flux and flux span from the FVA. The x-axis represents the analysis in context to the different host models; (B) Bubble plot of the rescued (hypothetically by using host resources - pink) or vital (green) metabolic reaction subsystems during infection in CAL. (C) Schematic workflow to investigate the metabolic alteration in CAL and host-Candida integrated model to identify the potential target for the analysis; (D) Plot of the normalized growth of CAL in individual and in an integrated model upon varying reaction bounds of Arg biosynthetic pathway reactions.

Although similarities were observed between murine models and cell lines, the analysis also highlighted some significant disparities. For instance, reactions for tryptophan and fatty acid metabolism became non-essential during CAL interactions with mammalian cell lines, but these reactions were essential for CAL survival in murine models. Interestingly the murine-based infection also showed an insignificant impact on the Arg metabolism of CAL. Besides the differences between the model systems (murine and cell lines), distinct metabolic adaptations between the cell lines were also identified. Variations in lipid metabolism were observed in CAL interactions with different cell lines; HUVEC cells showed enrichment in glycerophospholipid metabolism, while OKF6 cells exhibited enrichment in sphingolipid metabolism, highlighting cell line-specific metabolic adaptations (Figure 1A).

Essentiality analysis indicated that the host compensated for the requirements of tryptophan and fatty acid during infection, rescuing their essentiality. While improved essential reactions requirement in CAL infecting HUVEC, reactions related to terpenoid metabolism were rescued in OKF6 infection compared to control (Table S1, Figure 1B). Although multiple pathways of CAL appear altered in response to systemic infection, the significance of Arg biosynthesis during CAL infection was consistently implied based on functionally active pathways, enrichment of flux variations, flux span variations, and essential survival reactions. Consequently, genes involved in Arg biosynthesis emerged as promising targets for inhibiting CAL growth.

### 2.2. Integrated host-pathogen models reveal *ALT1* as a metabolic bottleneck

A first-of-its-kind comprehensive host-fungal metabolic model was developed by integrating the CAL metabolic model (*iRV781*)^22^ with the human metabolic model (Recon3D)^29^ to evaluate the metabolic dependency of CAL on the human host during infection. The schematic workflow employed to leverage the host-CAL integrated model for identifying potential targets that induce lethality in CAL is presented in Figure 1C. The host-CAL_WT_ model confirmed the growth of both CAL (140.17 mmol gdw^-1^ h^-1^) and the host (358.98 mmol gDW^-1^ h^-1^) under DMEM conditions. The exploratory analysis, including growth profiling and deletion studies (ref section 4.2.1), was performed on reactions of the Arg biosynthetic pathway (Table S5) on both the CAL_WT_ and host- CAL_WT_ models. The growth profiling showed that blocking the alanine transaminase (‘ALATA_Lm’) reaction that helps in the conversion of 2-oxoglutarate to glutamate while converting alanine to pyruvate halted the growth of CAL both in the presence and absence of host (Figure 1D). This effect was unique to ALATA_Lm, as other reactions like cytoplasmic and mitochondrial ‘ASPATh’ reaction, which converts 2-oxoglutarate to glutamate while converting aspartate to oxaloacetate did not significantly impact the growth of CAL under similar conditions. It can be assumed that the ‘ASPATh’ reaction is rescued when CAL infects and hijacks the host metabolic system for its survival.

A deeper investigation through FVA of the functionally mutant ALATA_Lm host-CAL_MT_ strain and the individual CAL_MT_ demonstrated that the deletion of ‘ALATA_Lm’ reaction significantly reduces the flux through reactions involved in glycerophospholipid metabolism in CAL_MT_ during infection (Table S7 and S8) without impacting the growth of host. Further comparative flux analysis of wildtype CAL (CAL_WT_) and host-CAL_WT_ and mutated (‘ALATA_Lm’ deleted) CAL (CAL_MT_) and host-CAL_MT_ revealed the depletion of glycerophospholipid metabolism in the host-CAL_WT_ in contrast to its enrichment in host-CAL_MT_. This could signify that the host metabolism cannot compensate for the glycerophospholipid production required in CAL, indicating that targeting the ALATA_Lm reaction (*ALT1*) could be an effective strategy for combating CAL infection.

### 2.3. *Candida* adapts to host stimuli by upregulating arginine synthesis

To evaluate the effect of host interaction in the expression of Arg metabolism genes of CAL, qPCR analysis of 41 genes belonging to this pathway was performed using RNA extracted from both CAL grown without (control) and with host vaginal A-431 cells (co-culture) for 5 h indicating early infection time-point^27^.

Arg metabolism can be subdivided into Arg biosynthesis and Arg and Pro metabolism (Figure 2). Arg biosynthesis involves a series of reactions that produce Arg from 2-oxoglutarate. Initially, 2-oxoglutarate is converted into glutamate (Glu) by alanine transaminase (*ALT1*) or aspartate aminotransferase isozymes (*AAT1/AAT21/AAT22*). Glu is then metabolized to Arg through two routes: (i) via induction of *ARG* genes, or (ii) through proline intermediates. In route 1, Glu is converted into ornithine by *ARG2*, *ARG5* and *ARG8*, which then enters the urea cycle and is converted into Arg. Route 2 follows the course of Arg production through Pro intermediates catalyzed by *GDH2*, *PRO1*, *PRO2* and *CAR2*.

**Figure 2.**
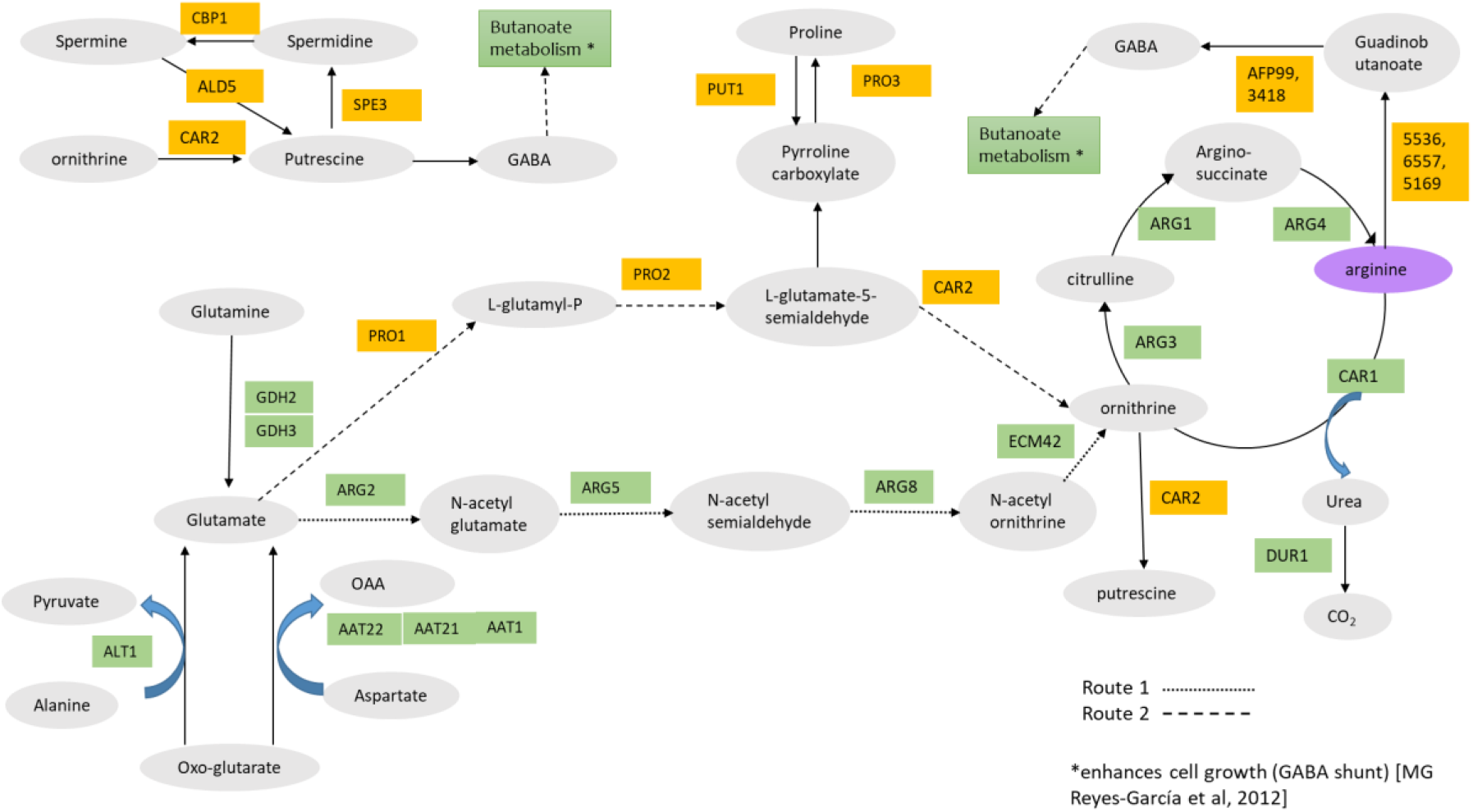
Pathway diagram of Arg metabolism based on data from KEGG pathway database. Metabolites are depicted in oval shapes, and enzymes are represented in rectangles; molecules involved in Arg biosynthesis (KEGG pathway cal00220) are shown in green, and those participating in Arg and Pro metabolism (KEGG pathway cal00330) are highlighted in yellow. Route 1 and Route 2 indicate the metabolic reactions involved in utilizing Glu for Arg synthesis.

After synthesis, *CAR1* converts Arg to ornithine and urea. Urea is then hydrolyzed to CO_2_ by the *DUR1,2* enzymes. These two reactions participating in the atrazine degradation pathway are important for CO_2_ induced hyphal morphogenesis in CAL^62^. Arg can also be metabolized into GABA by uncharacterized genes (*CaO19.5536, 6557, 5169*) and *AFP99*. Another route for GABA production is from spermine and putrescine catalyzed by products of *CBP1* and *SPE3*.

Under *in vitro* conditions, an overall upregulation was observed for genes belonging to the Arg biosynthesis and Arg and proline metabolism pathways (Figure 3A). While *CAR1* and *DUR 1,2* were upregulated both in the presence and absence of host cells, expression of genes *ARG3*, *ARG8,* and *ECM42* were downregulated in the presence of host cells. An upregulation was also observed for the genes *GDH2*, *PRO1*, *PRO2*, and *CAR2*, which are involved in the Arg synthesis from Glu through Pro intermediates (Figure 3B). Quantification of amino acids revealed a more pronounced increase in Arg compared to Glu and Ala, which are co-metabolized during Arg production, in CAL when grown with the host versus without the host. (Figure 3C). Together, these changes suggest that the host stimulates the production of Arg from Glu by activating the route 2 genes.

**Figure 3.**
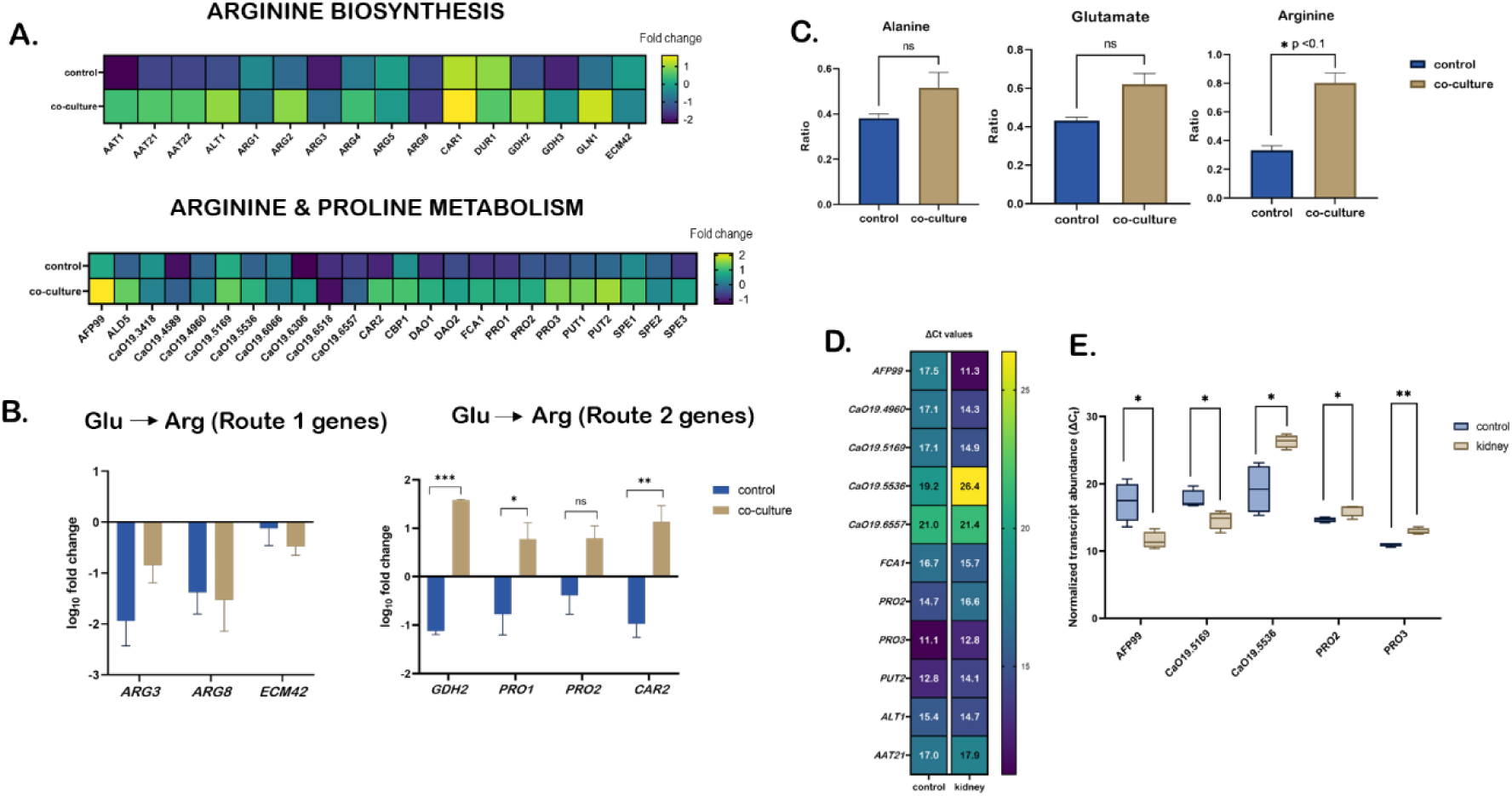
Impact of host-CAL interaction in Arg metabolism and evaluation of the eleven prioritised CAL target genes using systemic candidiasis murine model. (A) Heatmaps of expression of 41 genes involved in Arg metabolism in CAL (control) and in direct co-culture with vaginal A-431 cells at 5 hr time point. The color key represents the fold change in gene expression. (B) Bar plots depicting expression of genes involved in Arg synthesis from glutamate. Route 1 is through induction of ARG genes - ARG8, ARG3 and ECM42. Route 2 is through proline intermediates catalyzed by genes GDH2, PRO1, PRO2 and CAR2, which are upregulated during host interaction. (C) Comparison of intracellular levels of Ala, Glu and Arg in CAL when grown with and without host. Glu and Ala are intermediates in Arg synthesis. Ratio (y-axis) is obtained by dividing the concentration of amino acids of control or co-culture samples with the concentration of amino acid quantified from CAL cells used as inoculum for the experiment. (D) Expression of eleven prioritized targets of CAL from infected mice kidney. CAL used as inoculum for infection is taken as control. (E) Box and whisker plot illustrating the expression of five genes that exhibit significant differences in transcript level between control and infected mice kidney samples. No difference in transcript levels was observed for internal controls (18S) between groups (data not shown). All values are represented as mean ± SEM. Statistical significance was tested using unpaired t-test with Welch’s correction.

### 2.4. Prioritized targets in Arg metabolism exhibit variable *in vivo* expression patterns

Our data showed that the direct interaction of CAL with the host stimulates Arg synthesis by upregulating Arg metabolism. Next, to shortlist targets from the Arg metabolism pathway for extensive wet-lab validation, the 41 genes evaluated for expression levels through *in vitro* co-culture system (Figure 3A) were further screened using the in-house developed PBIT_v3_ algorithm^30^. This tool predicts pathogen targets based on essentiality, virulence, druggability, and non-homology to human and human gut microbiota. Of these screened proteins, 24 were identified as essential/virulent, and non-homologous to the human or gut microbiota proteome. While all 24 proteins were predicted to be druggable based on homology to known drug targets, 18 were found to be antigenic (see supplementary table S11). A subset of these genes (*ALT1*, *AAT21*, *CaO19.4960*, *AFP99*, *PRO2*, *PRO3*, *PUT2*, *FCA1*, *CaO19.5536*, *CaO19.6557*, and *CaO19.5169)* which were upregulated in host-CAL co-culture system (Figure 3A), were evaluated for *in vivo* expression during CAL infection in mice. Assessment of these genes in CAL-infected mice kidneys showed that for six genes, no significant difference was found in the transcript level as compared to control (Figure 3D). Of the remaining five genes, *CaO19.5536*, *PRO2* and *PRO3* transcripts were significantly depleted, while *AFP99* and *CaO19.5169* were significantly elevated in infected kidneys 48 h post-infection (Figure 3E). The gene *CaO19.5169* encodes a putative amidase; *in silico* druggability analysis showed that this protein and other two amidases (*CaO19.5536, CaO19.6557*) catalyzing the same reaction can be targeted by the drug 1-Dodecanol (1-DD), a fungistatic agent. When treated with 1-DD in YNB media at reported concentrations^31^, a decrease in the growth of CAL was also observed (Supplementary Figures 3D and 3E).

Among the shortlisted targets, *ALT1* and *AAT21* were identified as novel, although their *in vivo* expression was only marginally affected. *ALT1* exhibited a slight increase in expression compared to *AAT21* (Figure 3D). Findings from the integrated host-CAL model clearly indicated that disrupting the ALATA_Lm reaction catalyzed by *ALT1* significantly compromised the pathogen’s survival. In contrast, the reaction (ASPATh) catalyzed by *AAT21* did not have any such effect (Figure 1D). Based on this evidence, we sought to elucidate the role of *ALT1* in CAL virulence.

### 2.5. The *ALT1* knockout mutant displays attenuated virulence *in vitro*

To investigate its role in virulence, the *ALT1* deletion mutant (MT) was assessed for key virulence factors, including filamentation, secretion of aspartyl proteinases (SAPs), and phospholipase B activity. Furthermore, the interaction of the MT strain with the A-431 cell line was investigated *in vitro* using SEM, providing detailed structural insights into host-pathogen interactions.

The MT strain exhibited significantly reduced growth compared to the WT strain in YNB media supplemented with 2% glucose. After 24 h of growth, the MT strain showed a 66% reduction in dry weight (Supplementary Figure 1D). To induce the filamentous phenotype, both MT and WT strains were cultured in spider media at 37 °C for three days. The MT colonies appeared smooth with defined edges, while the WT colonies were wrinkled and exhibited peripheral invasive filaments (Figure 4A). Additionally, the MT strain produced significantly (p<0.01) less Arg than the WT strain (refer to Figure 5B).

**Figure 4.**
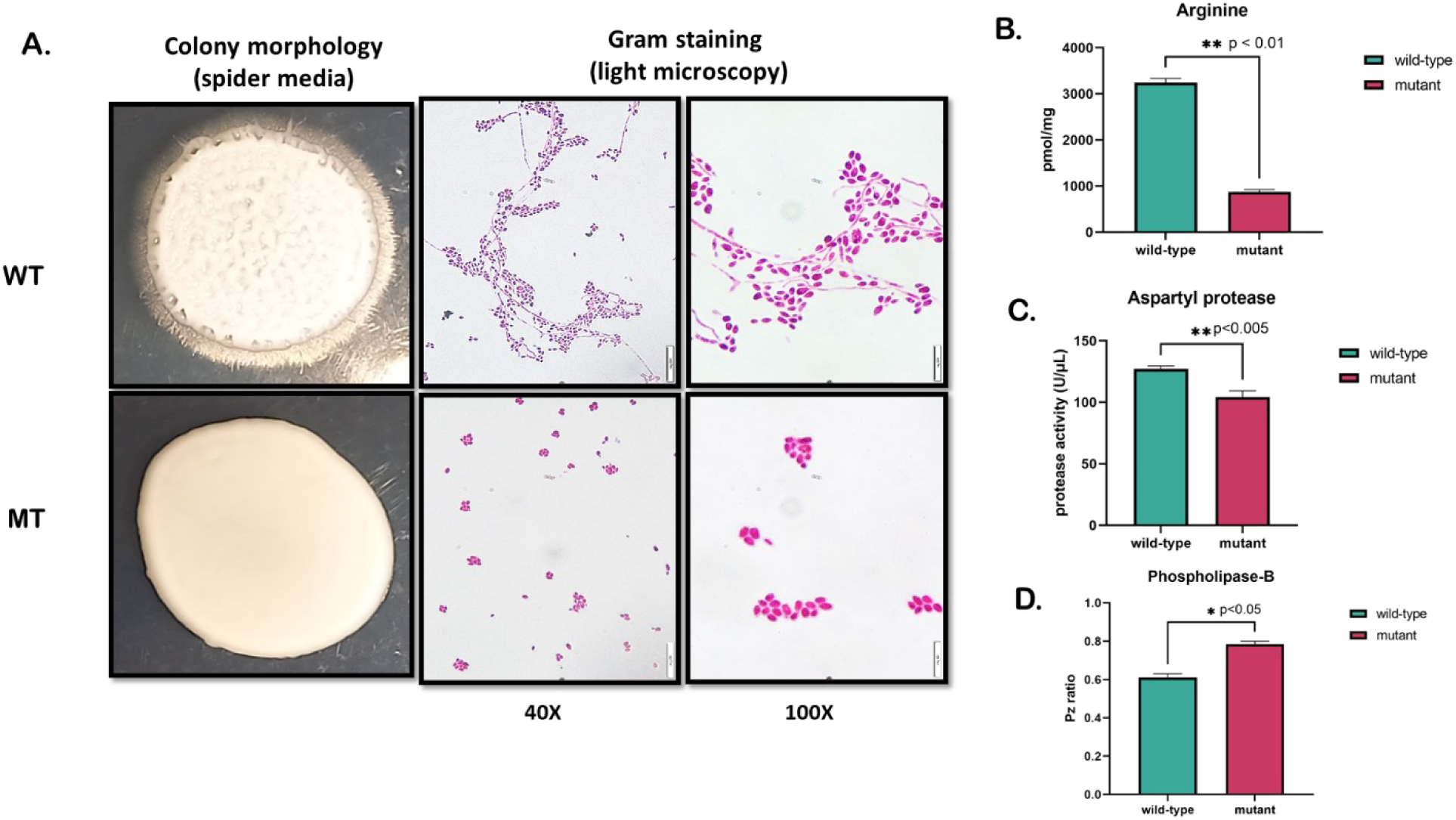
Comparison of the morphologies, intracellular Arg content and activity of host-damaging enzymes of wild-type (WT) and mutant (MT) CAL strains. (A) Colony characteristics of WT and MT strains after 72 h filamentation assay on spider media. WT and MT were stained with crystal violet and observed under 40X and 100X objectives. (B) Intracellular Arg content of WT and MT strains after 24 h growth in YPD media. (C) Aspartyl protease activity and (D) Phospholipase activity of WT and MT CAL strains. All values are represented as mean ± SEM. Statistical significance was tested using unpaired t-test.

**Figure 5.**
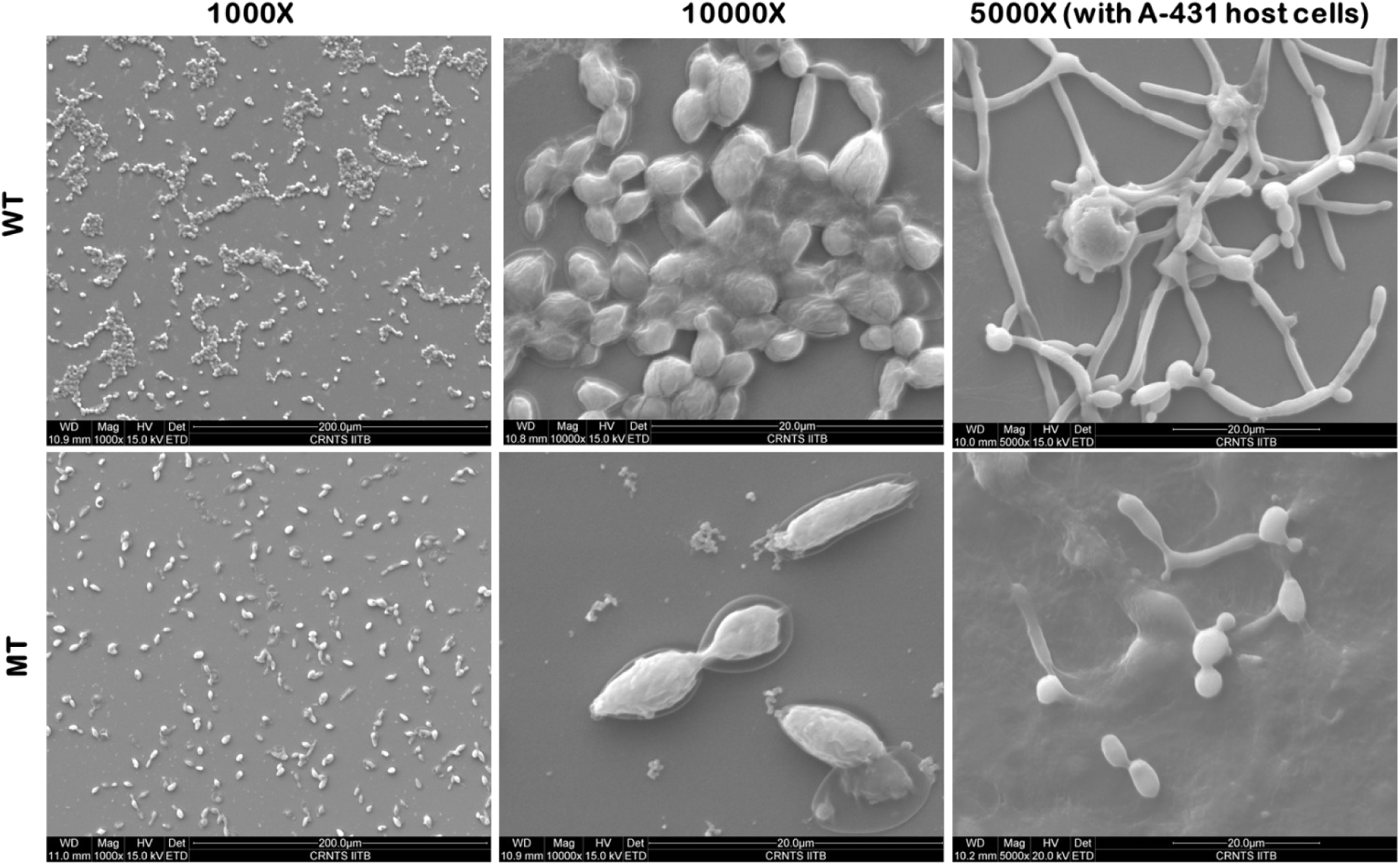
Scanning electron microscopy images of WT and MT singly and in co-culture with host cells.

To further confirm the absence of filamentation in the MT strain, both WT and MT cells were stained with crystal violet and observed microscopically. The MT cells lacked the hyphal morphotype characteristic of the WT colony cells (Figure 4A). Similarly, calcofluor white staining after 18 h of growth in YNB media revealed no hyphal or pseudohyphal formation in MT strain (Supplementary Figure 2). The loss of filamentation in the MT strain was further verified by SEM analysis, where MT and WT were grown in DMEM for 5 h (Figure 5). MT cells exhibited a spindle-shaped morphology, contrasting with the round shape of WT cells. A noticeable reduction in extracellular matrix (ECM) was observed in MT, possibly contributing to the loss of cell-to-cell adherence and colonization. In contrast, WT formed clusters with extensive ECM deposition (Figure 5). However, when MT was co-cultured directly with host cells, a short hyphal phenotype was observed. While the WT strain showed abundant and elongated hyphal structures, the MT hyphae were notably shorter and less developed in comparison.

The activity of host-damaging enzymes, specifically aspartyl proteases and phospholipases, was quantified using their respective assays. Both SAP and phospholipase activity were reduced in the MT strain. A significant decrease in SAP activity (p < 0.005) was observed in MT compared to WT after 24 h of growth in YNB (Figure 4C). Phospholipase activity was also significantly lower (p<0.05) in MT (Pz = 0.77), while the WT strain exhibited moderate activity (Pz = 0.63) (Figure 4D).

### 2.6. CAL *ALT1* is essential for *in vivo* pathogenicity

*In vivo* pathogenicity of the MT strain was assessed based on its ability to induce systemic candidiasis in a murine model over a 15-day survival study. Male C57BL/6 mice were infected with 1 x 10^5^ – 1 x 10^6^ MT and WT cells, and their survival was monitored for 15 days. No mortality was observed in the group injected with MT strain (n=10) throughout the study duration. However, animals in the group injected with WT strain (n = 10) experienced mortality on days 6 (n=1), 8 (n=2), 9 (n=2), and 14 (n=1) (Figure 6A). Animals infected with the WT strain exhibited notable weight loss, whereas no significant weight changes were recorded in the group infected with the MT strain (Figure 6A).

**Figure 6.**
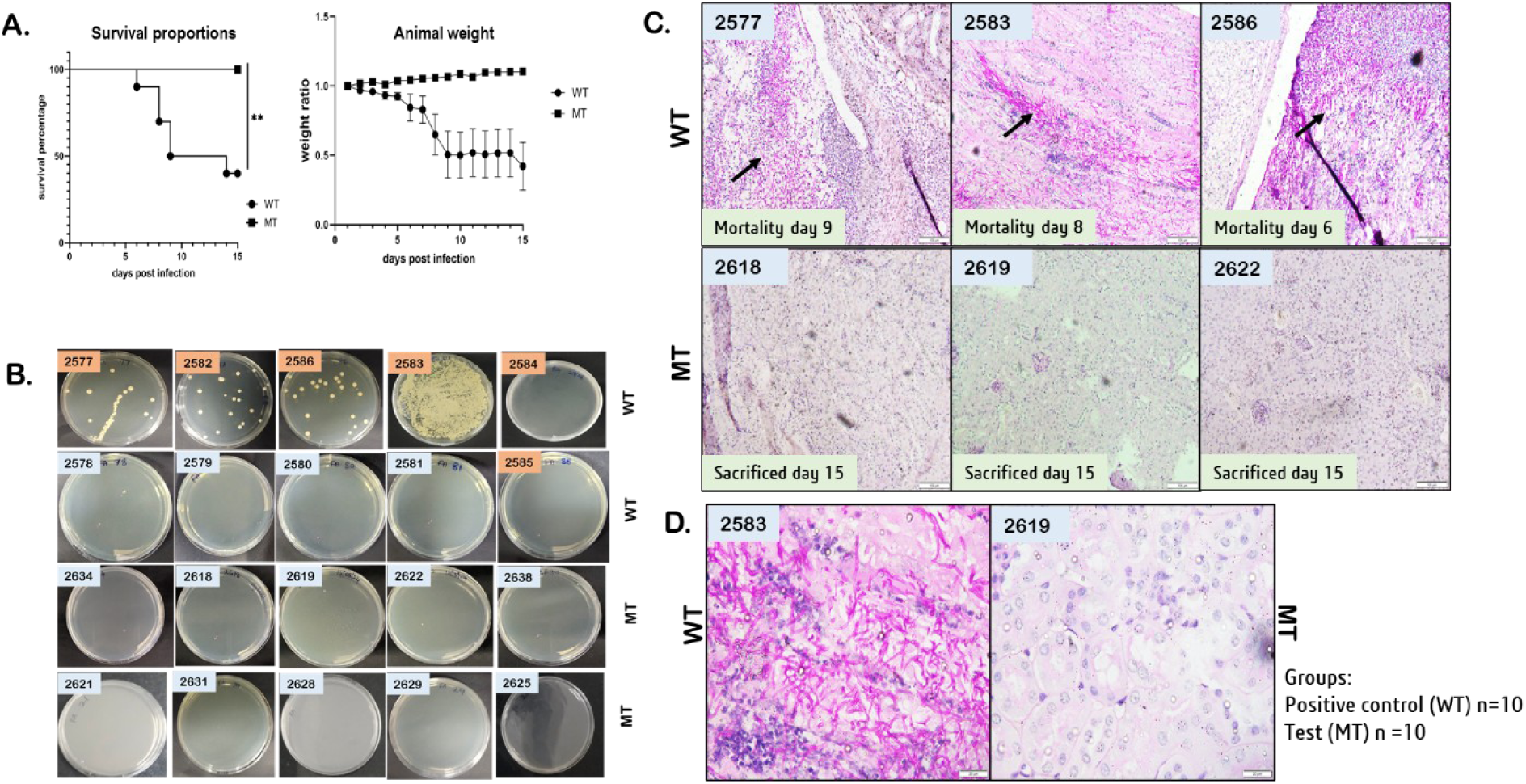
Evaluation of virulence of MT CAL alt1Δ (MT) in C57BL/6 mice. (A) Survival plot of mice following the administration of WT (n =10) and MT (n = 10) CAL strains. Statistical analysis was performed using log-rank (Mantel-Cox) test using Prism 8.0 (**P < 0.005). Fluctuations in body weight of WT and MT infected groups as observed during the study. As compared to the MT-infected group, a decrease in body weight (indicative of candidiasis) was observed for mice injected with wild-type. (B) Determination of fungal burden using fungal agar plates. The animal identification number is depicted in rectangular boxes upper-left corner of images. Blue boxes indicate animals that survived and orange boxes indicate animals that died during the study period. (C) Representative images of kidney sections of WT and MT infected groups under 10X magnification. Arrows indicate the presence of fungal hyphae, (D) Representative images of kidney sections from WT and MT groups under 40X magnification. Mice infected with WT show the presence of fungal hyphae (in pink) and invading neutrophils (in blue) in the kidney sections. Animal identification numbers are indicated in blue boxes at the upper-left corner of images.

To quantify fungal burden, kidney homogenates from infected mice were plated on YPD agar containing chloramphenicol (100 μg/mL) and incubated for 48 hours. At the end of the 15-day study, no fungal burden was detected in the kidneys of MT-infected mice. In contrast, mice that succumbed to WT infection exhibited high fungal load characterized by the presence of fungal hyphae and invading neutrophils in the kidney sections (Figure 6C-D). Interestingly, no fungal burden was observed in the kidneys of surviving WT-infected mice (n=5) sacrificed on day 15 (Figure 6B).

## 3. Discussion

Our integrative approach, combining systems biology metabolic modeling with experimental validation, has presented valuable insights into the metabolic alterations in CAL during host infection and identified potential therapeutic targets.

We leveraged the potential of transcriptomics data and GSMM of CAL (*iRV781*) to investigate the metabolic rerouting in CAL during host infection. Although these models reflected only minimal variation in the number of reactions (Figure 1A), a significant variation in reaction fluxes between the enriched and repressed states was observed. For example, flux through reactions of steroid and terpenoid backbone synthesis pathway in CAL under infected state was enriched in HUVEC while reduced in OKF6 (Figure 1A), suggesting environment-specific metabolic modulation. Notably, while the maximum flux remained constant in steroid, sphingolipid metabolism and terpenoid backbone synthesis across conditions (as evidenced by insignificant flux variation, Figure 1A), the observed differences in flux span indicate a potential metabolic redundancy within CAL. This redundancy likely represents an adaptive strategy, allowing the pathogen to maintain metabolic flexibility in diverse host niches. This, in addition to variation in the number of essential reactions between HUVEC and OKF6, depicts the distinct internal metabolic wiring (varying functional pathways) at different environmental states (control and infection) in different cell lines (Table S1). The loss of essentiality of fatty acid biosynthesis and metabolism, tryptophan metabolism, phenylalanine, tyrosine and tryptophan metabolism in CAL during infection suggests that CAL relies on host metabolic support for its growth and survival. This metabolic hijacking is consistent with the prior observations on other pathogens, such as increased lipid accumulation during *M. tuberculosis* infection and the redirection of fatty acid metabolism away from glycolysis in *S. typhimurium* infection^32–34^. It is interesting to note that, unlike pathways like fatty acid biosynthesis, steroid biosynthesis, carbon metabolism, and biosynthesis of other amino acids, the significance of arginine biosynthesis was consistently emphasized in all our analyses, as illustrated in Figures 1A and 1B. The inability of host metabolism to compensate for the impact of arginine biosynthesis pathways in CAL summons a more in-depth investigation into its function during host-pathogen interaction.

To gain deeper insights into the metabolic interplay between CAL and host in Arg biosynthesis during infection, flux profiling of thirteen unique reactions of arginine biosynthesis pathway in host-CAL_WT_ and CAL_WT_ was conducted. As observed from Figure 1D, limiting the flux through the reactions ALATA_Lm, ASPATh, ASPAThm, and GALh negatively affected the growth rate of CAL_WT_. However, this limitation was relieved in the presence of the host (integrated host- CAL_WT_ model) for all reactions except for ALATA_Lm catalyzed by the alanine transaminase (*ALT1*) enzyme. The negative impact on CAL growth while limiting ALATA_Lm highlighted the critical role of metabolites derived from *ALT1* reaction for CAL growth and maintenance, which cannot be compensated by cross-feeding nutrients from the host. Comparative flux analysis of wild type (CAL_WT_) and mutant (CAL_MT,_ ALATA_Lm deleted CAL) with host (host-CAL_WT_ and host-CAL_MT_) exhibited the depletion of carbon substrate (fructose and mannose), glycerophospholipid metabolism and reduction in amino acid biosynthesis in the host-CAL_WT_ and enrichment of glycerophospholipid metabolism in host-CAL_MT_ (Table S7 and S8). This implies that CAL_WT_ depends on the metabolic resources of the host to optimize its survival during infection. In contrast, the mutant strain (CAL_MT_), unable to hijack host metabolism, must rely on its own metabolic pathways to meet its metabolic needs. Thus, the activation of glycerophospholipid metabolism was possibly a compensatory mechanism to adapt to the stress caused by disrupted arginine biosynthesis. Similar adaptations have been documented in *Saccharomyces cerevisiae* under hypoxic conditions, where glycerophospholipid metabolism plays a crucial role in maintaining membrane stability and cellular homeostasis^35^. The investigative analysis of host-CAL models assures that the disruption in Arg biosynthesis by *ALT1* in CAL_MT_ might have triggered the glycerophospholipid activation to improve its pathogenicity in the host (Table S8).

The implication of Arg biosynthesis has been documented through various pieces of evidence. In our previous study, we identified 12 metabolic targets in *C. tropicalis*, the majority of which were associated with the Arg biosynthesis pathway^24^. Similarly, Jiménez-López et al. reported that CAL upregulates Arg biosynthesis genes *ARG1* and *ARG3* in response to reactive oxygen species (ROS) within macrophages^36^. Furthermore, Ghosh et al. observed that L-Arg promotes CAL growth and filamentation, facilitating its escape from murine macrophages^37^. Nonetheless, a comprehensive investigation of all arginine biosynthesis genes to identify therapeutic targets has not been previously conducted.

A gene expression study was conducted on all CAL genes involved in arginine biosynthesis, as well as in arginine and proline metabolism, using an *in vitro* host-CAL model system. In the presence of host cells, a net upregulation of genes of this pathway and a significant increase in Arg production (Figure 3A-C) was observed. We hypothesize, that Arg production in the pathogen is most likely skewed toward route 2 of arginine biosynthesis pathway via proline intermediates (Figure 3B), a trend also highlighted by Jiménez-López et al.^36^ when CAL was subjected to ROS-induced stress. Among these route 2 genes, *GDH2* has been previously linked to mitochondrial function during macrophage interaction^38^, while others (*PRO1, PRO2*, and *CAR2*) were regulated by protein kinase Gcn2/Gcn4 as a response to amino acid starvation^39^. This suggests that Arg production is essential during co-culture, likely due to the stress imposed by nutrient competition between fungal and host cells, to promote hyphal growth for better nutrient scavenging^40^.

Unconventional metabolic pathways present new opportunities to combat drug resistance in *Candida*^41^. Our research identified arginine metabolism as a key pathway essential for host-pathogen dynamics, suggesting its potential for alternative therapeutic development. Selected genes within this pathway, shortlisted through the PBIT_v3_ pipeline as potential drug/vaccine candidates, were subjected to further investigation in a murine systemic candidiasis model. Significant differences in expression were observed only for two genes (AFP99 and CaO19.5169), 48h post infection (Figure 3E). These genes convert Arg to GABA (Gamma-aminobutyric acid). In CAL, GABA has been linked to various functions such as hyphal development, promotion of phospholipase activity^42^, and SAP production through modulation of PKA and MAPK pathways^43^. Thus, the antifungal activity of 1-DD, as seen in Supplementary Figure 2, may be due to the suppression of GABA production by inhibition of CaO19.5169 and allied amidases.

Although there was marginal upregulation of the novel *ALT1* gene *in vivo* (Figure 5A), its central role in amino acid sensing and metabolism (Figure 7), along with findings from *in silico* analysis (Figure 1D) and *in vitro* gene expression (Figure 3A), prompted us to speculate the significant impact of *ALT1* knockout on *Candida* virulence. For validation, the *ALT1* gene deletion mutant (MT) was created, and as expected, it showed a significant decrease in internal Arg content compared to WT based on the results of amino acid analysis by HR-LCMS (Figure 4B). Further, *in vitro* and *in vivo* assays suggested that MT exhibited characteristics of an avirulent/attenuated strain, such as poor growth, reduced hyphal formation, decreased extracellular matrix production, and lowered synthesis of host-damaging enzymes (Figures 4 and 5). However, during co-culture with host cells, MT could produce a shorter hyphal form (Figure 5). This hyphal induction may be a compensatory response triggered by the nutrient exchanges between host and fungal cells, as suggested by an increase in flux in the transporters pathway, as depicted in the systems biology analysis (Table S8). *In vivo* virulence studies in a mouse model showed that animals infected with MT exhibited 100% survival compared to <50% survival in the WT-injected group at the end of day 15 (Figure 6). It is possible that the loss of virulence characteristics in MT led to faster clearance of the injected pathogen by the host immune system; this may have implications for vaccine development and is subjected to further investigation. Contradictorily, the surviving mice in the WT group also cleared the infection, likely due to a heightened immune response, consistent with previous findings in C57BL/6 mice during epidermal infections^44^. These findings indicate that the MT strain is incapable of initiating an infection, thereby establishing the role of *ALT1 in vivo* CAL pathogenicity and virulence.

**Figure 7.**
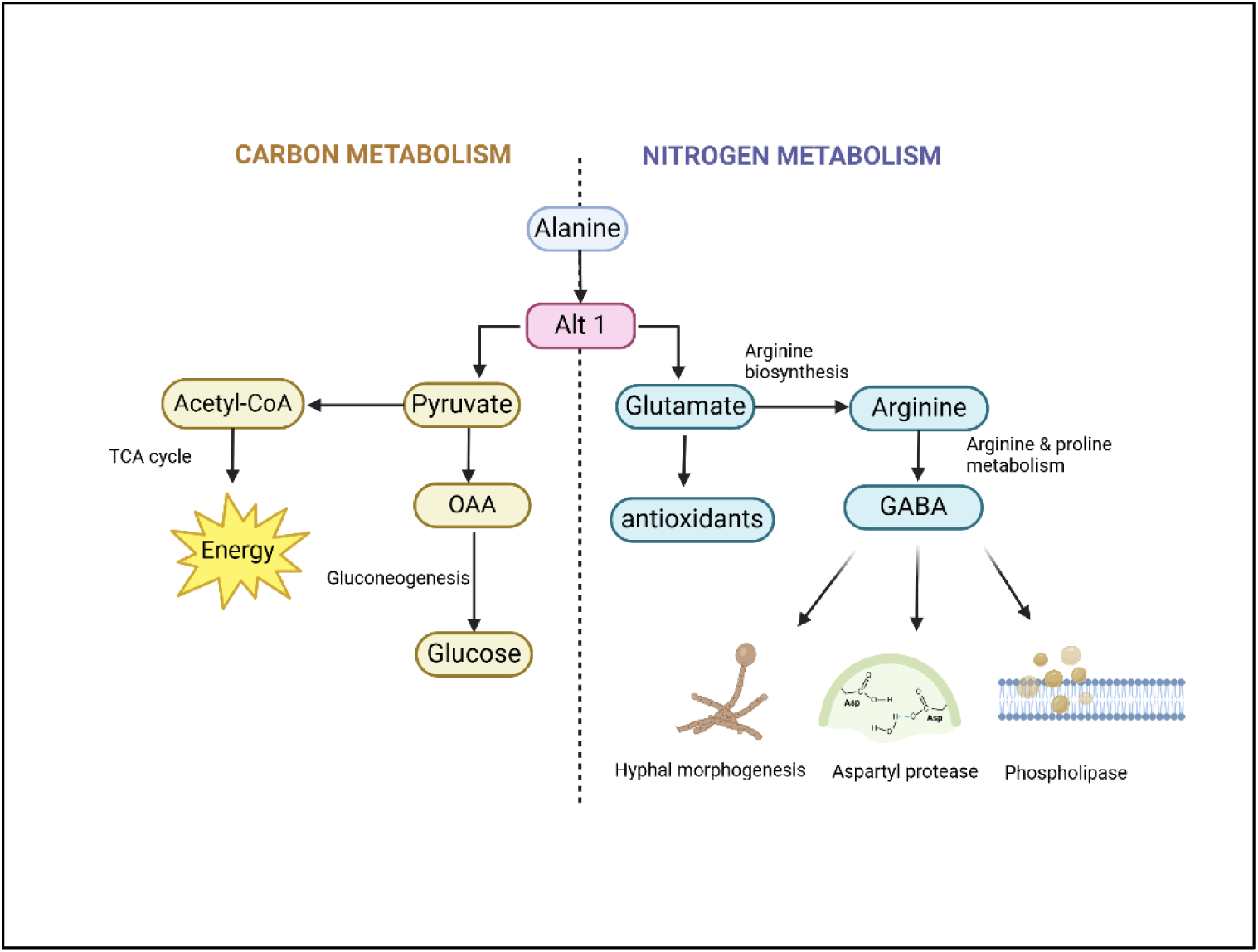
Role of ALT1 in CAL virulence. Regulation of carbon and nitrogen metabolism by ALT1. Pyruvate formed during the reaction is used for energy generation in the TCA cycle and for the production of glucose by gluconeogenesis. Glu participates in Arg biosynthesis, which eventually affects the GABA shunt regulating hyphal formation, hydrolyzing enzymes and also the production of antioxidants.

In CAL, the gene (ca*ALT1*) is predicted to be linked with mitochondrial function based on its homology to *Saccharomyces cerevisiae* sce*ALT1* and is annotated as a putative alanine transaminase enzyme in UniProt^45^ and the Candida Genome Database^46^. In *S. cerevisiae*, a decrease in Arg leads to repression of GABA shunt genes *GAD1*, *UGA1*, and *UGA2* along with impaired mitochondrial DNA integrity and gene expression^47^. Additionally, deletion of *sceALT1* has been shown to negatively affect the chronological lifespan of *S. cerevisiae*^48^. Building on these insights and our findings, we speculate that *ALT1* disruption in *C. albicans* may reduce Arg availability, thereby impairing GABA production. Given the functional role of GABA in CAL virulence, this reduction could have significant implications for mitochondrial function and pathogen fitness. Since *ALT1* is critical for CAL virulence and its deletion renders the pathogen avirulent, this attenuated strain can be decisively leveraged as a live attenuated vaccine or utilized as a strategic target for novel antifungal treatments.

Our study does have some limitations. Our predictions hinge on the accuracy of the GSMMs available for CAL and the human host. The current GSMMs (*iRV781*) are semi-curated, incomplete, and require additional data to precisely represent metabolic parameters at any given time point. Nevertheless, these models have been shown to be very potent in developing a deeper understanding of pathogen metabolism^49^, and enable the prediction of drug targets^50^. The human Recon3D model also simplifies the host to a single cell, rather than a complex organism with diverse tissues and cell types. Yet, since we have experimentally validated some insights derived from the host-CAL metabolic model, we have shown that these models can undoubtedly provide valuable insights into infections and host-pathogen interactions.

Overall, we present three main contributions from our study. For the first time, a host-CAL integrated metabolic model was developed and validated by *in vitro* and *in vivo* experiments. Second, we established that during host-CAL interaction and infection, there is distinct metabolic rewiring in CAL; these findings could be exploited for novel therapeutic strategies. Third, we showed that arginine biosynthesis plays a pivotal role in the growth and maintenance of CAL during infection and its essentiality cannot be compensated by the host. Further, the metabolite depletion caused by the deletion of the *ALT1* catalyzing reaction could not be compensated by host nutrients, and the depletion of Arg through deletion of the *ALT1* gene attenuates CAL virulence. Taken together, these observations make a compelling case for targeting Arg biosynthesis.

In summary, our study underscores the potential of targeting metabolic vulnerabilities in fungal pathogens for therapeutic purposes. The validated host-CAL model developed herein can provide profound insights into host-CAL interactions and aid in predicting novel therapeutic strategies. By integrating computational metabolic modeling with *in vitro* cell line-based and *in vivo* murine-based models, the *ALT1*-deleted CAL strain emerges as a promising vaccine candidate for candidiasis. Additionally, it paves the way for *ALT1* inhibitors as promising antifungal therapeutics and supports systems biology as a platform for novel target discovery in pathogens.

## 4. Materials and methods

### 4.1. Construction of context-specific CAL models

#### 4.1.1. Transcriptomics data processing

The transcriptomic profile of CAL with mammalian cell lines and murine models using GSMM was investigated to evaluate the metabolic alterations in CAL during infection. Transcriptomics data from healthy mammalian cell lines (HUVEC - Human Umbilical Vein Endothelial Cells and OKF6 - Oral Keratinocytes) and those infected with CAL strain SC5314 were retrieved from the Gene Expression Omnibus database (GEO accession ID: GSE56093^27^). The data for healthy and infected cell lines at the 0th, 5th, and 8th hours were examined to study the temporal dynamics of *Candida* infection. Flux analyses were performed to identify shared and cell line-specific coping mechanisms during *in vitro* infection. The transcriptional profile of systemic CAL infection in mice at the 24^th^ hour was also included to examine the events of infection in the mammalian host and pathogen (GEO accession ID: GSE83682^28^).

#### 4.1.2. Reconstruction of context-specific Candida models

The processed transcriptomics data of uninfected CAL and CAL, following infection of mammalian cell lines (GSE56093) and murine hosts (GSE83682), were incorporated into the GSMM of CAL (*iRV781*) to simulate the metabolic state of CAL cells under normal and infection conditions. The *iRV781* model represents the metabolic structure of CAL with 1,221 reactions, 927 metabolites, and 781 genes^22^. Since the gene annotations differed from those in the transcriptomics data, they were modified by mapping the annotations to the Candida Genome Database (CGD)^46^ to achieve a consensus annotation between the model and transcriptomics data. To integrate the gene expression data, we employed GIMME (Gene Inactivity Moderated by Metabolism and Expression)^51^, an objective-based transcriptomic data integration algorithm. A threshold of 10% median gene expression was used to filter out the genes or reactions that should be excluded from the model. The context-specific CAL models were then evaluated for growth in YPD media. All metabolic modeling simulations of CAL and host-CAL models were conducted using CobraToolbox v3^52^ and MATLAB R2022a.

### 4.2. Systematic analysis of metabolic alteration during CAL infection

Initially, the reactions in the context-specific CAL models under infected and healthy/uninfected conditions were extracted to compare the active reactions during infection. Active reactions refer to those included in the model based on gene expression data; the genes associated with these reactions have expression levels above the threshold, effectively representing the expressed metabolic genes.

#### 4.2.1. Analysis of metabolic enrichment during infection

The metabolic capacity of context-specific metabolic models of CAL, generated for both uninfected and infected conditions, was analyzed using Flux Variability Analysis (FVA) as described by Mahadevan and Schilling^53^. FVA calculates the minimum and maximum possible fluxes for each reaction while ensuring the overall metabolic balance of the model. Reactions exhibiting a difference of more/less than 20% in maximum flux or flux span (difference of maximum to minimum flux) between uninfected and infected conditions were identified as key contributors to the metabolic adjustments of CAL in response to the host environment. While the maximum flux could hint at the reactions with changes in maximum potential, the flux span approach might identify reactions showing tightened or relaxed fluxes with the same maximum flux, which could imply the other possible metabolic adaptation in the microorganism.

#### 4.2.2. Essentiality analysis during infection

Essentiality analysis was performed on the context-specific CAL models by simulating the deletion of each gene or reaction. Reactions (or genes) were classified as essential if the growth rate of the knockout strain differed by more than 10% compared to the wild-type strain. In the analysis, reactions were labelled as *’rescued’* if they were essential only in the uninfected condition, and *’needed’* if they were shown to be essential only during infection.

#### 4.2.3. Flux enrichment analysis of the selected reactions

Flux enrichment analysis (FEA) was performed on the reactions identified through FVA and essentiality analysis to prioritize the significantly impacted and critical metabolic pathways during CAL infection. The Cobra toolbox suite was utilized for FEA, employing a hypergeometric test with an adjusted p-value < 0.05.

### 4.3. Construction and analysis of host-CAL metabolic ,odel

#### 4.3.1. Generation of the integrated host-CAL metabolic model

An integrated host-CAL model was constructed to investigate the metabolic dependencies between the human host and CAL during infection. Recon 3D model^29^ and *iRV781*^22^ were used to represent the human host and CAL, respectively. The reaction and metabolites of the CAL model were renamed to ensure its compatibility with the host model. The altered CAL model was integrated into the host model by creating a common metabolite exchange compartment (‘[u]’) to share the resources between the host and CAL. The integration was facilitated by the ‘*createMultipleSpeciesModel*’ functionality in CobraToolbox v2, followed by the coupling of intracellular and extracellular reactions within the combined host-CAL model to their respective biomass components (Figure 1C). For instance, CAL reactions were coupled with CAL biomass, and host reactions were coupled with host biomass and biomass maintenance reactions. The integrated model was subjected to growth assessment on the provided medium, wherein the model was optimised for the growth sum of both the host and CAL.

#### 4.3.2. Analysis of the host-CAL integrated model under knock-out conditions

The growth of CAL in the reconstructed host-CAL model was monitored for varying fluxes of the reactions involved in Arg biosynthesis in CAL. The growth profile of both CAL_WT_ and integrated host-CAL_WT_ was evaluated for the reaction lower bound ranges of [-10, 50] for all the thirteen reactions in the Arg biosynthesis pathway. After the genes/reactions had been shortlisted for the analysis, the mutant strain (deletion of ‘ALATA_Lm’) was created by setting the corresponding lower bounds to 1e-5 (Step-2 in Figure 1C). These mutant models (CAL_MT_ and host-CAL_MT_) were generated using CAL_WT_ and host-CAL_WT_ metabolic models, respectively, as a base model. Both the models were subjected to FVA, followed by the examination of reactions with varying fluxes. The significant flux changes between the wild-type and mutant models were selected if the ratio of wild type and mutant flux is greater/lesser than 20%. The significantly altered reactions were analysed for three different sets of models (Set-1: host-CAL_WT_ vs host- CAL_MT_; Set-2: CAL_WT_ vs host-CAL_WT_; Set-3: CAL_MT_ and host-CALMT) to understand the metabolic reprogramming in CAL during systemic candidiasis. The identified reactions were subjected to FEA to identify the metabolic subsystems that are significantly altered during host infection.

### 4.4. *In silico* prediction of potential targets

Protein sequences of all genes (n=41) belonging to the Arg metabolism pathway were subjected to comparative proteomics analysis using the in-house developed tool PBIT_v3_^30^(https://www.pbit.bicnirrh.res.in/) for prioritizing therapeutic targets. Non-homology analysis was performed against the proteome of human and gut microbiota using sequence identity < 50% and e-value < 0.0001. Non-homologous proteins were then screened for their essentiality, virulence, druggability, and antigenicity; a sequence alignment length cut-off of 1% and e-value < 0.0001 were used for the analysis.

### 4.5. Expression analysis of genes of prioritized pathway using infection models

#### 4.5.1. In vitro co-culture model of vaginal candidiasis

A-431 human vaginal epithelial cells were seeded in 6-well plates at a density of 3 × 10^5^ cells per well in Dulbecco’s Modified Eagle Medium (DMEM) supplemented with 10% fetal bovine serum (FBS). Cells were incubated overnight at 37 °C in a humidified atmosphere containing 5% CO₂ to reach approximately 90% confluency. CAL SC5314 (ATCC MYA-2876) was cultured overnight at 37°C in yeast extract peptone dextrose (YPD) medium. The culture was harvested by centrifugation at 4000 rpm for 5 min, washed twice with sterile phosphate-buffered saline (PBS), and resuspended in serum-free DMEM. The CAL inoculum was adjusted to a concentration of 3 × 10^6^ cells/mL. The prepared inoculum was added to each well containing the A-431 monolayer. Co-cultures were incubated for 5 h at 37 °C in serum-free DMEM. In parallel, an aliquot of the CAL inoculum was used as a time-zero control for gene expression normalization. Control wells containing CAL alone (without host cells) were also incubated under the same conditions.

#### 4.5.2. In vivo model of systemic candidiasis

Male C57BL/6 mice (6-8 weeks old, n=5) were used to establish a systemic candidiasis model. The mice were housed under controlled conditions with a temperature of 23 ± 1°C, humidity of 55 ± 5%, and a 14 h light/10 h dark cycle, with unrestricted access to food and water. Each mouse was injected with 1 x 10⁶ yeast cells of CAL strain SC5314 via the lateral tail vein, while the remaining inoculum was used as a negative control.

After 48 h post-infection, the mice were humanely euthanized, and their kidneys were collected and stored at -80 °C till further analysis. The experimental procedures were approved by the Institutional Animal Ethics Committee (IAEC), following the guidelines of the Committee for the Purpose of Control and Supervision of Experiments on Animals (CPCSEA; project no: 17/21).

#### 4.5.3. RNA extraction and qPCR analysis

RNA extraction from the *in vitro* vaginal candidiasis model was performed by harvesting co-cultured cells. After removing the supernatant, cells from each well were scraped into 500 µL of chilled TRIzol reagent (Ambion, Life Technologies), snap-frozen in liquid nitrogen, and stored at -80 °C until further processing. The experiment was conducted in triplicate, with each technical replicate generated by pooling two wells. For RNA extraction, frozen samples were thawed on ice, and 500 µL of AE lysis buffer^54^ (50 mM sodium acetate, pH 5.3, 10 mM EDTA, 1% SDS) was added to each tube. Samples were gently vortexed and then subjected to mechanical lysis using bead-beating procedure. Lysing matrix G (MP Biomedicals, USA), containing 1.6 mm silicon carbide particles and 2 mm glass beads, was used at a speed of 6 m/s for 30 seconds in the FastPrep-24™ instrument (MP Biomedicals, USA) for lysing CAL cells.

RNA extraction from the CAL-infected kidneys of the systemic candidiasis model was performed as follows: Frozen kidneys were thawed on ice and homogenized in TRIzol reagent: AE lysis buffer (1:1) using a mechanical homogenizer at 8000 rpm on ice. The homogenized tissue was further processed using bead beating with lysing matrix G, as described previously.

The resulting cell and tissue lysates were centrifuged at >12,000 × g, and the supernatant was used for RNA extraction. RNA isolation was performed using the phenol:chloroform:isoamyl alcohol (25:24:1, v/v) method. RNA was precipitated overnight at -20 °C in isopropanol. cDNA was synthesized from 2 μg of RNA using the SuperScript™ III First-Strand Synthesis System (Thermo Fisher Scientific). Real-time PCR was performed using CAL-specific primers (Supplementary table S9) and SYBR green master mix (BioRad). For the *in vivo* model, gene expression was analyzed only for eleven genes - *ALT1*, *AAT21*, *CaO19.4960*, *AFP99*, *PRO2*, *PRO3*, *PUT2*, *FCA1*, *CaO19.5536*, *CaO19.6557*, and *CaO19.5169*, that were found to be upregulated in the *in vitro* model and predicted as therapeutic targets through *in silico* analysis (Table S11). Relative gene expression was calculated using the ΔΔCt method, and 18s rRNA was used as an internal reference.

### 4.6. Determination of amino acid content in CAL during host interaction

CAL and A-431 cells were co-cultured following the protocol detailed in Section 2.5.1. After a 5 h incubation, the supernatant was discarded, and the cells were scraped into chilled PBS. The cell suspension was then centrifuged, and the resulting pellet was stored at -80 °C for future processing. Before storage, human cells were selectively removed from the co-cultured samples by treating the cell pellet with a buffer containing 4 M guanidine thiocyanate, 25 mM sodium citrate, 0.5% Sarkosyl (N-lauroyl-sarcosine), and 0.1 M β-mercaptoethanol, which selectively lyses human cells^55^.

For sample preparation, the stored cell pellets were thawed on ice and air-dried, and 100 mg of the sample was hydrolyzed in 10 mL of 6 N HCl at 110 °C for 24 h. A 20 μL aliquot from the hydrolysate was evaporated using a speed vacuum and reconstituted in 50 μL of 0.1 N HCl. Amino acids involved in the Arg metabolism pathway – Ala, Arg and Glu, were measured using high-resolution LC-MS (HR-LCMS) analysis conducted at the SAIF facility, IIT Bombay.

### 4.7. Construction of deletion mutant of alanine transaminase (*ALT1*) gene in CAL

Deletion of the selected target gene, alanine transaminase (*ALT1*), was carried out using the SAT1 flipper cassette as described by O. Reuß et al.^56^, following the approval of the Institute Biosafety Committee (IBSC) for project 1/2020. The sequence of the *ALT1* gene (orf19.346, chromosome 3), along with 1,000 bp upstream and downstream, was retrieved from the Candida Genome Database^46^ for primer design (see Supplementary File S2). The upstream fragment, 5′ flank F1 (0.38 kb), was custom synthesized by Abgenex (Bhubaneshwar, India) and cloned into the pUC57 plasmid between the *ApaI* and *XhoI* restriction sites.

The downstream fragment, 3′ flank F2 (0.42 kb), was amplified via PCR from CAL genomic DNA using forward (F2_F) and reverse (F2_R) primers containing *NotI* and *SacI* restriction sites, respectively. After digestion with the appropriate restriction enzymes, both fragments were ligated using NEB quick ligase to the upstream and downstream of SAT1 flipper sites of plasmid pSFS2A, which encodes the nourseothricin resistance marker. The constructed plasmid pSFS2A_*ALT1*, containing the 5 kb gene deletion cassette for *ALT1*, was propagated in *E. coli* DH5α competent cells (HiMedia) following transformation.

As outlined in supplementary figure 1A, the 5 kb gene deletion cassette was excised from pSFS2A_*ALT1* via restriction digestion using Quickcut ApaI and Quickcut SacI (Takara Inc., Japan). The cassette was then introduced into CAL by transformation using PEG/LiAc reagents in the Yeastmaker™ Yeast Transformation System 2 kit, following the modified protocol of Walther and Wendland^57^. Transformed colonies were selected on YPD agar plates (YPD medium with 15 g/L agar) containing antibiotics nourseothricin (200 µg/mL) after incubation at 30 °C for 3-4 days. The transformation was repeated twice to obtain homozygous mutant colonies. Successful gene deletion was confirmed through colony PCR with gene-specific primers.

### 4.8. Quantification of virulence parameters of *ALT1* deficient CAL

#### 4.8.1. Filamentation assay and comparison of morphology using light microscopy

A filamentation assay was performed, and colony morphologies were compared between wild-type (WT) and *ALT1* mutant (MT) strains. A 10 µL suspension of both strains (1 × 10^6^ cells/mL) in YPD (Yeast Peptone Dextrose) was spotted onto solid Spider medium (composed of 1% nutrient broth, 1% mannitol, and 0.2% K_2_HPO_4_). The plates were incubated at 37 °C for 3 days. Colony morphology was observed, and individual colonies were collected, smeared onto clean glass slides, and heat-fixed. The smears were then stained with crystal violet to examine the cell morphology under a light microscope.

#### 4.8.2. Scanning electron microscopy (SEM) in the presence of host cells

The interaction of WT and MT strains with vaginal epithelial host cells (A-431) was analyzed using SEM. Both strains (1 × 10³ CFU) were cultured on polylysine-coated coverslips for 5 h in DMEM medium without FBS, either in the presence or absence of host cells. After incubation, the coverslips were washed with sterile PBS and fixed overnight at 4 °C using 0.3% glutaraldehyde. The following day, the coverslips were sequentially dehydrated in graded ethanol (30% to 70%) and air-dried. They were then sputter-coated with platinum and examined using SEM (Quanta 200) at SAIF, IIT-Bombay.

#### 4.8.3. Secreted aspartyl protease assay

Supernatants from overnight cultures of WT and MT strains grown in YNB media were collected by centrifugation at 3000 g for 5 min. A 50 µL aliquot of the supernatant was mixed with 150 µL of 0.1 M sodium citrate buffer (pH 2.8) and 150 µL of 1% (wt/vol) azocasein, followed by incubation for 1 h at 37 °C. 200 µL of 20% trichloroacetic acid was then added to terminate protease activity and precipitate undigested proteins. The mixture was then centrifuged again at 3000 g for 5 min. The resulting 400 µL supernatant was combined with an equal volume of 500 mM NaOH and incubated at 25°C for 15 min. Absorbance was measured at 440 nm using a spectrophotometer to assess protease activity. Protease activity (U/µL) was calculated using the formula from Benmebarek et al.^58^:

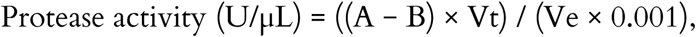

where A and B represent the optical densities of the negative control and the WT/mutant samples, respectively; Vt is the total reaction volume, and Ve is the volume of enzyme supernatant.

#### 4.8.4. Phospholipase B assay

A 10 µL suspension of WT and MT strains in PBS, containing 10^6^ cells/mL, was spotted in the petri plate containing egg yolk medium (Himedia) and allowed to air-dry at room temperature. The plates were incubated at 37 °C for 72 h. Phospholipase activity, measured by the Pz index, was considered positive when a distinct precipitation zone appeared around the colonies. The level of phospholipase production was determined by calculating the ratio of the total diameter (colony plus precipitation zone) to the diameter of the colony alone. Pz values between 0.999 and 0.700 indicated low enzymatic activity, values from 0.699 to 0.400 indicated moderate activity, and values between 0.399 and 0.100 signified high enzymatic activity^59^.

### 4.9. Assessment of *in vivo* pathogenicity and virulence of *ALT1* deficient CAL

The study to assess *in vivo* pathogenicity of *ALT1*-deficient CAL (MT) was conducted following approval from the Institutional Animal Ethics Committee (IAEC) at ICMR-NIRRCH (Project No: 7/24). The experiments were carried out under previously specified controlled conditions.

Male C57BL/6 mice, 8-10 weeks old and weighing 25-30 g (n=20), were randomly assigned to two groups: (i) a positive control group (n=10) injected with the WT CAL strain, and (ii) a test group (n=10) injected with *ALT1*-deficient MT strain. Each mouse received a weight-adjusted inoculum of 1 x 10⁵ to 1 x 10⁶ colony-forming units (CFU) via intravenous injection into the lateral tail vein to initiate systemic infection^60^.

The mice were monitored daily for changes in body weight and survival over a 15-day period. Survival analysis was performed using the Kaplan-Meier method^61^, wherein a death event was assigned a value of 1, and survival was assigned a value of 0. Weight ratios were determined by dividing the mice’s body weight each day by their initial body weight recorded on day 1. CAL load in harvested kidneys was estimated by agar plate method and histologically after Periodic Acid Schiff (PAS) staining of kidney sections. Data analysis was conducted using GraphPad Prism 8.

### 4.10. Statistics and reproducibility

All experiments were replicated at least twice. For experiments, mean and standard error (SD) are reported in the figure legends for technical replicates from representative experiments performed in duplicates or triplicates. Statistical significance was determined as indicated in the figure legends.

## Code and data availability

All data are available within the article and its supplementary materials provided along with the manuscript. The codes to reproduce the systems biology analysis are available on GitHub: https://github.com/Indupalani/Host-Candida-metabolic-modelling/tree/main

## Acknowledgements

The authors SIT, SC, and YG are grateful to Dr. Geetanjali Sachdeva, Director, ICMR-NIRRCH for support. This work (RA/1803/12-2024) was supported by was supported by DBT, India [BT/PR40165/BTIS/137/12/2021], SERB, India [CRG/2021/004937] and SRF from ICMR [Myco/Fell/14/2022-ECD-II]. IP acknowledges the funding received by PMRF, MOE, India. The authors IP and KR acknowledge the financial support from IBSE and WSAI, IIT Madras, India.

## Author contributions

SIT, KR and KVV participated in acquisition of funding, project administration, and supervision. SIT, SC, KR, and IP participated in the research design. KVV provided concept design and guidance for the study. SC, IP and KR designed systems biology studies; IP executed systems biology analysis. SIT and SC designed wet lab experiments; SC performed *in vitro* and animal experiments. YG provided technical support in animal experiments. SC and IP wrote the first draft of the manuscript. SIT and KR critically reviewed the manuscript. All authors read and approved the final version of the manuscript.

## Abbreviations

CAL: Candida albicans
NACs: Non-albicans Candida species
GSMM: Genome scale metabolic model
ALT1: Alanine transaminase
CAL_WT_: Wild type CAL metabolic model
CAL_MT_: ALT1 deleted CAL metabolic model
Host-CAL: Host CAL integrated model
WT: wild-type CAL strain
MT: ALT1 deleted mutant CAL strain
ALATA_Lm: L alanine transaminase mitochondrial reaction
ASPATh: Aspartate aminotransferase reaction

