## Supplementary file_S1 for "An integrated systems biology approach establishes arginine biosynthesis as a metabolic weakness in *Candida albicans* during host infection"

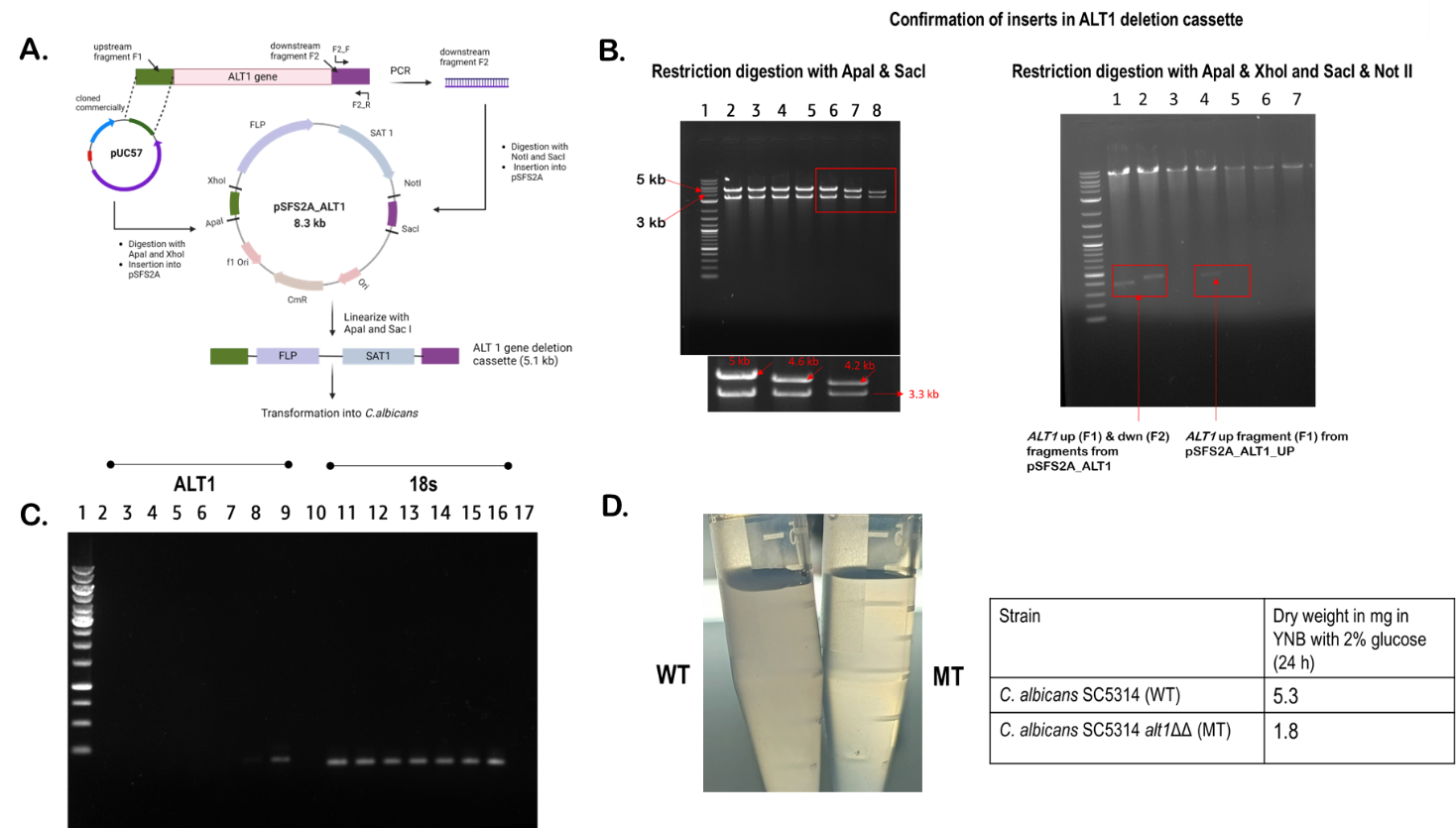

***Supplementary figure 1****. (A) Construction of ALT1 deletion cassette using pSFS2A plasmid for homologous recombination-based gene deletion in CAL. (B) Restriction digestion (RD) map of plasmid pSFS2A_ALT1 with deletion cassette (lane 2-6), Plasmid integrated with* upstream *fragment F1 of ALT1 gene (lane 7) and parent plasmid pSFS2A without any inserts. The second figure shows RD of plasmids pSFS2A_ALT1(wells 1-3), pSFS2A_ALT1_up (lane 4-5) & pSFS2A (lane 6-7) with ApaI & XhoI (lane 1, 4 & 6) and SacI & NotI (lane 2, 5 & 7). (C) PCR to check the disruption of ALT1 gene in CAL. Lane 2-7: PCR products of transformed colonies of CAL with ALT1 specific primers, lane 8: Positive control from WT CAL colony, lane 9 & 17: NTC for ALT1 & 18S, lane 10-15: PCR products of transformed colonies of CAL with 18S specific primers, and lane 16: Positive control from WT CAL colony. (D) Cell density and dry cell weight of WT and MT CAL grown in YNB media after 24 h.*

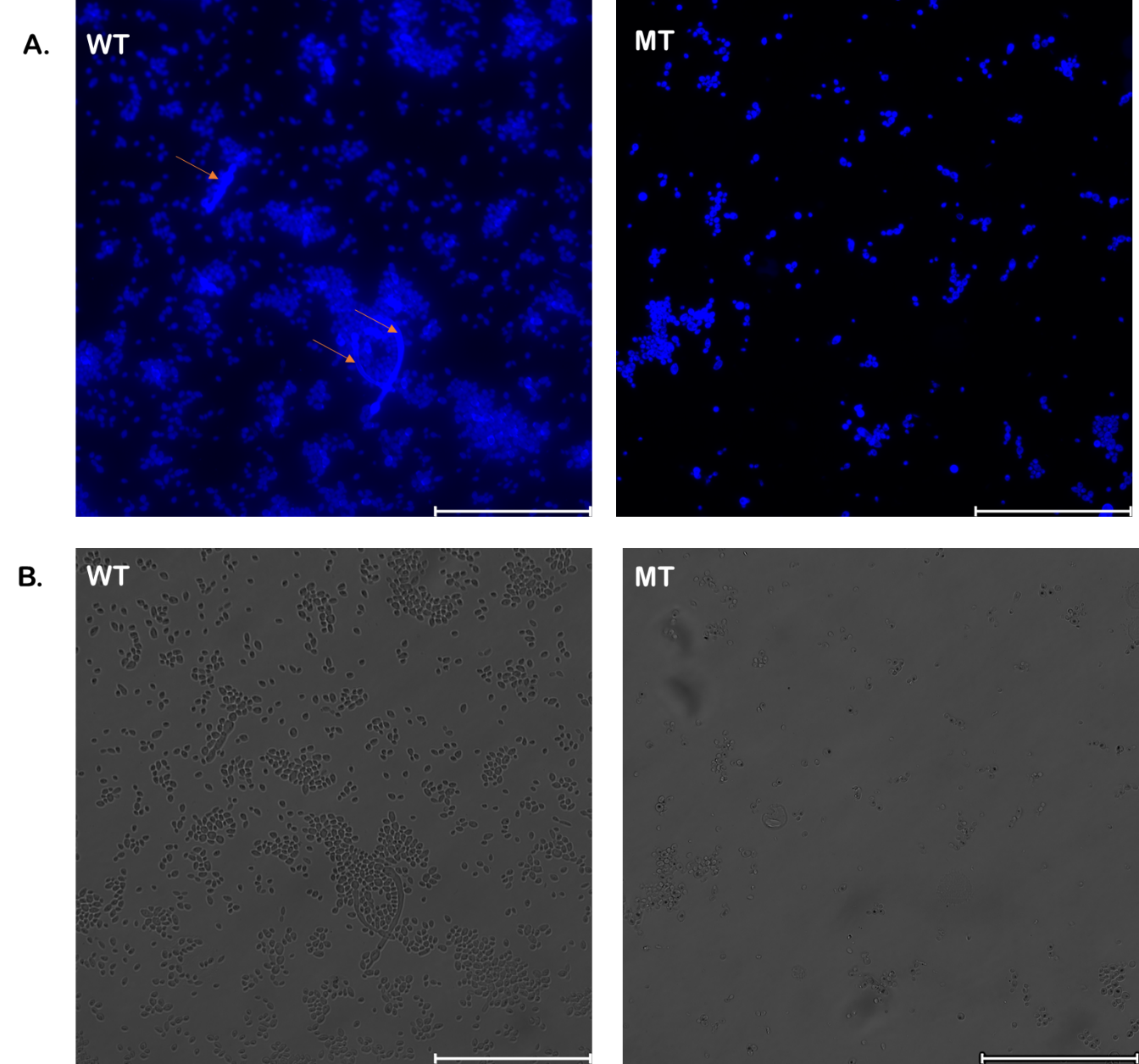

**Supplementary figure 2.** *(A) Fluorescent and (B) phase microscopic images of wild-type (WT) and mutant (MT) CAL after 18 hr growth in hypha-inducing YNB media. Hyphae are marked with red arrows*

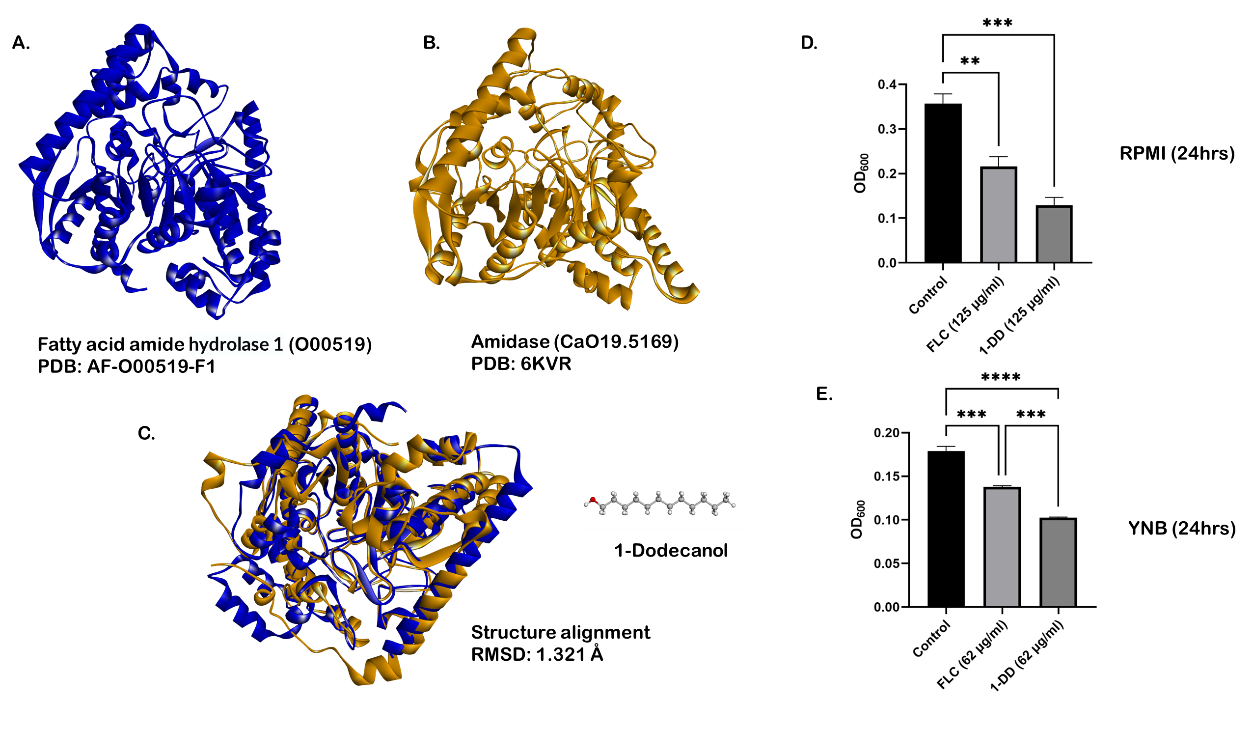

**Supplementary figure 3.** *Structural homology between targets of 1-Dodecanol. (A) Fatty-acid amide hydrolase [UniProt ID: O00519, PDB:* *AF_AFO00519F1 (in blue), and (B) orf19.5169 (putative amidase, UniProt ID: G1UA68, PDB: 6KVR) (in gold). (C) Structure alignment (using BIOVIA Discovery Studio 2021) shows an RMSD of 1.31Å. (D) Drug assays using 1-Dodecanol (1-DD) in RPMI media, and (E) YNB media expressed as optical density (OD) values at 600 nm.*

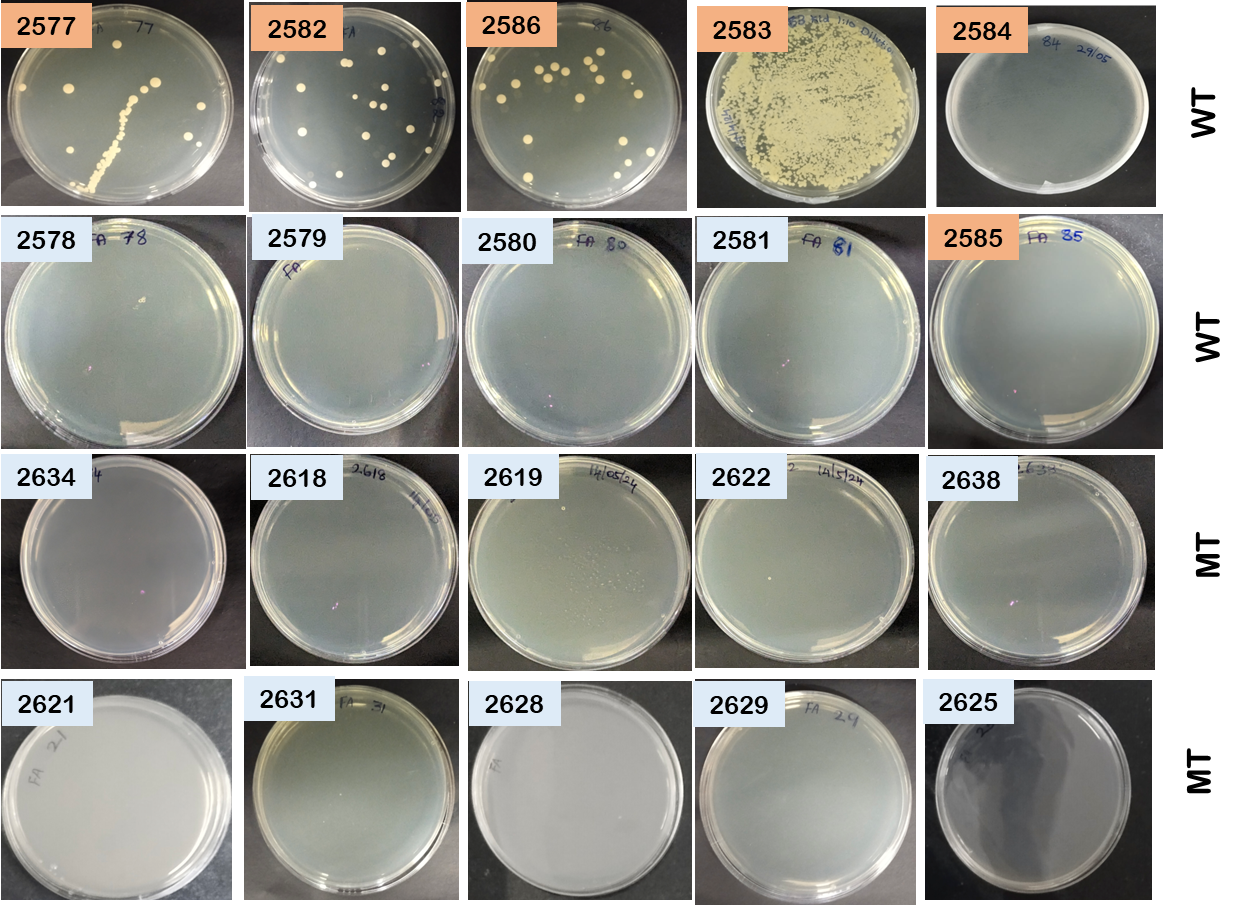

***Supplementary figure 4****. Determination of fungal burden using fungal agar plates. The animal identification number is depicted in rectangular boxes upper-left corner of images. Blue boxes – animals that survived during the study, orange boxes – animals that experienced mortality during the study period.*
