## Supplementary file_S2 for "An integrated systems biology approach establishes arginine biosynthesis as a metabolic weakness in *Candida albicans* during host infection"

Candida albicans ALT1 gene sequence ± 1kb (start and end site marked in bold)

>C3_03480C_A COORDS:Ca22chr3A_C_albicans_SC5314:738106-741668C (3563 nucleotides)

ATAAACAAAAACTGTTAATAACTTTATAGTTAACTTTAAAGCCGCTAGCGGCTCGAAGTTTTGCTGGCATCTTTAGTTTCCACGGCAAAAATTAAAACCCAAGAAAGAGCGGTATTCAAAGTTTAGATAATAGTTTTGCATCACATTAATTAGGTTATCTTGTTTTTATTCAAATAGTCATTCTCTCCACCGCTTAAAAACAATTCGAACGTCGGACATAAACTTCCGAAATATCCACCTAATGATGTTGGAAACACCGATCACAAACCAAGCTTATCAATGTTTAATTGGCAACGTGGACAAATGAGTTTAAAGATTCAAATTTACTATGTGGTTGATTGATTGAAGCATTTCTCAATCACCCAGATACAACATACTGAAGATAATTTAAGAAATATATAACAATTTAGTCTATAAGTAGCTATCCCTTTATTATCTAACAAATAGAAGTCACAAGATCATTAAGATGGGACCAAAAAAAATAGAAATCTATTTTGTCTTGTTTATTATCTTTTAGAAAGTTTTCGTGTAATTTTCTCCACTTTTTAATTTTTCTGACTCACCCACTCAACTCACTCCTAATCTACCCTTATCAATTTTCAACAACAACACACTCATATAATTCTTAATTCAGATTTGTTCTGCTATATATTATTCATTATTTTTTTTTTTCACCCCAAAGAACCCACAAAGAACCCACAAATAAAATCGAATTAATTTTCACAATTTTTTTCCCCTTTCTTTCTTTTCTTTCTTTTCAACTGATTTATTTCATTTCATTTCATTCTTTTGATTTACAACCAACTTTTCATTCACTTATTTATTGTTAGTATTGTGTTTCTCCTCCCTTCCCACTGTGTCACGTTATTTATAATACTATACTACCCTTTCCCCCGAACACAATAATAATCACAGGCATTTGAAACAAACAAAGAAACTAGATACTAATACATATATACAAACAAGAAAATAAATAAAAAAAAAAATTAGACAATCAATA**ATG**CTTAGAGTAAATTCACGATTATTAACTACTCGAAAATCTCAATTTATGTTCACTTCTAATTCCTTAAGGTTTTTAGCAACCTTTAAACCTGCGGATCCATTGACTACTCATGATATCAACCCACAAACGGTCGAGGCAAAATACGCTGTACGTGGGAAAATCCCAATTATTGCCGATGAATTGAATGAATTGATTCAAAAGCAACCACAGCTGCATGGATTACCATTTAGCAAAATCATTAATGCCAACATTGGTAACCCACAGCAATTGGAACAACGTCCATTGACATGGTATCGTCAAGTATTGTCTCTTTTACAATACCCGGATTTATTGAAAAATGGAGACCCTGAAACCGTTAAATCACTTTATCCTGAAGATGTGATTGAGCGAGCACAATCAATTTTGAAACACATTGGATCAATAGGGGCATACTCTCATTCTCAGGGTGCCAGTTATTTCCGACAATCTATTGCTGAATTTATAACTAACCGTGATGGTGGTTATGTATCCCACGCCAACAACATTTTTTTAACTTCCGGGGCATCAACTGCAGTGTCGTATTTATTACAAATTTTGTCTGTCAATGAAAACTCCGGGTTCTTAATTCCGATCCCACAGTATCCTTTGTATACTGCCACTATTGCCTTGAATAATGCCAAGCCAATTGGTTATTATCTCGATGAGTCGAACCATTGGTCAACCAATCCTCAAGAGATTAGAGAATTGATCGAAACCAATCAACTGCAAGGTATTAACATCAAGGCATTGGTGGTGATTAACCCGGGGAACCCAACGGGGGCAATTTTATCATCACAAGATATAATTGAATTGATAGATATTGCTGCTGAATATGGAATTGTATTAATTGCCGATGAGGTTTATCAAGAAAATATCTTCAAAGGGAAATTTGTTTCATTCAAGAAGATTTTGTCGGAATTAATTGAGCAAGACCCTCAAACTTATAAACATGTTCAATTAGCATCATTACACTCGACATCGAAAGGTGTTAGTGGAGAATGTGGACAACGTGGTGGATATATGGAATTAGTTGGGTTCAAACCGGAAGTTAAAGATGTGGTTTTCAAATTGGCATCGATTAATTTATGTTCTGTTGTCTCTGGTCAAGCATTAATGGAATTAATGATTAATCCTCCTCAAGAAGGTGATCCTAGTTACCCCTTGTACAAAAGCGAAACCGAGTCAATCCATAACGATTTGGAATCAAGAGCAGAATCACTCTATCAAGCATTTTTACAAATGGAAGATATCAAATGTAATAAACCAATGGGGGCCATGTATATTTTCCCCACTTTAGATTTTGATCCAGCCCAATACCATAAGTTATATTCGAGAGCTAAGAATTCCAATTTACAAATTGATGATATTTATTGTATTGAGTTATTAGAAGGTACAGGTATTTGTTGTGTTCCTGGTAATGGATTTGGTCAGAAACCAAATACTTATCATTTAAGAACAACATTTTTACCACCAGGTAAAGAATGGATTGATAAATGGATAAATTTCCATAAATCATTTATTAAAAAATATAAAGATGAA**TAG**ATACCCTATATATATTTATATATGTACATATATGTCTTGTCTATAAAATACAAAATATGATAACCAAAAAAAGAAAAAAAAAAAAATCAAAGTTAAGACACACATTCATTCAAAGTAGTTTCAGTGGAAACGGGAAGTTTAGTTATAGACATTACCTAAAGTAATTGATTTAATAACGGTATAGTCATCCATTCCATAAAGTGAACCTTCTCTACCAAAACCTGATTCTTTAACCCCACCAAATGGCAATGCAGCATCGGTGAATAATCCAGTATTAACGGAAACCATACCATTTTCCAAAAATTCTGACATATACCATACAGTATTCAAATTTTCCGAAAAGATATATGATGCCAACCCATAAGGAGTATCATTACACCATTGCAAAACTTGTTCTTTAGAATCAAATGGGATAATAGCTGCCAAGGGACCAAAAGTTTCTTCTTTAACAACCTTCATATCTTGAGTCACATCCTTAACTACTGAAGGGGAATAGAAATTTTCTCCCAATTGGGGCAATCTACCTCCTTCAACAATCAATTTTGCTCCTTTTTCAACGGCATCTTGAACATGATCTTCAACTTTTTCAATGGCTTTAGTGTTTATCAAACAACCATGAGTTACACCTGGTTCAAATCCATTACCAATTTTGAATTGATTTACTTTCTCAACAAATTTATTACAAAATTCATCATAAACCCCCTTTTCAACATAAATACGATTGGCACAAACACAAGTTTGCCCTAACGATCTGAATTTAGAAGTAATACTTTGATCAACAGCAAGATCTAAATTACAATCATTGAAAACAATAATCGGAGCATTACCACCTAATTCCATTGATAATTTTTTCAAAGTTGAAGATGATTGTTGCATTAATAATTTACCAACATTAGTTGATCCAGTGAAACTGATTTTTTTCAATTTTGGTGATTGACAAAATTTCAATCCACACATTGGAGTATTGGTTACAGAAGTCAATACAACATTGAAACAACCA

**Primer F2_F -** Forward downstream primer to amplify F2 sequence of *ALT1* for cloning using NotI [Tm:   62.7 °C]

GAGAAAGCGGCCGCCCAATGGGGGCCATGTATATT

**Primer F2_R -** Reverse downstream primer to amplify DS sequence of *ALT1* for cloning using SacI [Tm: 60.5 °C]

GAGAGGCGAGCTCCTTCCCGTTTCCACTGAAACTA

Sequence of upstream fragment F1

GGGCC**↑**CCAATTCGAACGTCGGACATAAACTTCCGAAATATCCACCTAATGATGTTGGAAACACCGATCACAAACCAAGCTTATCAATGTTTAATTGGCAACGTGGACAAATGAGTTTAAAGATTCAAATTTACTATGTGGTTGATTGATTGAAGCATTTCTCAATCACCCAGATACAACATACTGAAGATAATTTAAGAAATATATAACAATTTAGTCTATAAGTAGCTATCCCTTTATTATCTAACAAATAGAAGTCACAAGATCATTAAGATGGGACCAAAAAAAATAGAAATCTATTTTGTCTTGTTTATTATCTTTTAGAAAGTTTTCGTGTA ATTTTCTCCACTTTTTAATTTTTCTGACTCACCCACTCAACTCACTCC C**↓**TCGAG 3’

**↑ ApaI site**

**↓ XhoI site**

Sequence of downstream fragment F2

GAGAAAGC**↑**GGCCGCCCAATGGGGGCCATGTATATTTTCCCCACTTTAGATTTTGATCCAGCCCAATACCATAAGTTATATTCGAGAGCTAAGAATTCCAATTTACAAATTGATGATATTTATTGTATTGAGTTATTAGAAGGTACAGGTATTTGTTGTGTTCCTGGTAATGGATTTGGTCAGAAACCAAATACTTATCATTTAAGAACAACATTTTTACCACCAGGTAAAGAATGGATTGATAAATGGATAAATTTCCATAAATCATTTATTAAAAAATATAAAGATGAATAGATACCCTATATATATTTATATATGTACATATATGTCTTGTCTATAAAATACAAAATATGATAACCAAAAAAAGAAAAAAAAAAAAATCAAAGTTAAGACACACATTCATTCAAAGTAGTTTCAGTGGAAACGGGAAGGAGCT**↓**CGCCTCTC

**↑ NotI site**

**↓ SacI site**

Primers for verification:

**Primer ALT1_F**	CAAACCGGAAGTTAAAGATG

**Primer ALT1_R**	ATTCTGCTCTTGATTCCAAA
